# Capturing ribosomal structures in cellular extracts with cryoPRISM: A purification-free cryoEM approach reveals novel structural states

**DOI:** 10.1101/2025.08.21.669550

**Authors:** Mira B. May, Gabriella S. Lopez-Perez, Joseph H. Davis

## Abstract

Structural analyses of ribosomes by single particle cryogenic electron microscopy (cryoEM) have traditionally relied on purified or reconstituted samples, with particles often trapped in desired states using genetic, pharmacological, or biochemical perturbations. While informative, such *in vitro* methods often fail to capture the full diversity of structural states and associated protein factors present in cells. In contrast, *in situ* cryoelectron tomography preserves cellular context but is limited by low throughput and modest resolution. Here, we present cryoPRISM (<u>p</u>urification-free ribosome imaging from <u>s</u>ubcellular <u>m</u>ixtures), a rapid *ex vivo* workflow encompassing cell lysis, vitrification, and image analysis methods for high-resolution analyses of ribosomal structures directly from cell lysates. Applying cryoPRISM in *E. coli*, we resolved more than twenty distinct ribosomal states spanning assembly, translation initiation, elongation, trans-translation, and quiescence, including a novel configuration of EF-G bound to idle ribosomes with the ribosome hibernation factor RaiA. Given its speed, accessibility, and ability to preserve native interactions and structural heterogeneity, we anticipate that cryoPRISM will be broadly applicable for uncovering ribosomal biology across diverse organisms and conditions.

## INTRODUCTION

Protein synthesis is executed by the ribosome, whose assembly and activity are tightly regulated within its cellular environment. Across its life cycle, protein factors guide the ribosome as it transitions through a variety of compositionally and conformationally distinct states. Initially, ribosome biogenesis is orchestrated by a diverse set of assembly factors, including GTPases, RNA-modifying enzymes, and RNA helicases that are thought to assist in RNA folding, protein incorporation, and structural remodeling; often operating at specific stages of the assembly process (Shajani *et al*. 2011; Davis and Williamson 2017). Once mature, global conformational rearrangements in the ribosomal subunits are coupled with the activities of translational factors and transfer RNAs (tRNAs) to support protein synthesis (Korostelev 2022). For example, during translation initiation, protein factors IF-1, IF-3, and IF-2 bind to the small subunit and help ensure initiator tRNA pairs with the mRNA start codon (Gualerzi and Pon 2015). Subsequently, the large subunit is recruited, and the ribosome begins a translation elongation cycle where tRNAs and mRNA are shuttled through the mRNA decoding and peptidyl transferase centers, with the help of GTPases EF-Tu and EF-G (Voorhees and Ramakrishnan 2013).

Beyond canonical translation, ribosomes can adopt a variety of critical but often rare structural states that modulate their activity. Indeed, when mRNAs are truncated or otherwise damaged such that translating ribosomes stall, multiple pathways exist for bacteria to perform “ribosome rescue” (Müller *et al*. 2021; Ray and Apirion 1979). One such pathway, termed “trans-translation”, acts by exchanging truncated mRNA with a new open reading frame encoded on a transfer-messenger RNA (tmRNA), resuming translation from the tmRNA and tagging the peptide for degradation. When exposed to conditions that disfavor growth, bacteria respond by temporarily inactivating ribosomes as part of translational control (Maki and Yoshida 2022), wherein they can either form 100S disomes, facilitated by HPF and RMF (Wada *et al*. 1990; Yoshida *et al*. 2002; Polikanov *et al*. 2012), or they can remain as translationally silenced 70S monomers (Maki *et al*. 2000; Ueta *et al*. 2005). The latter is governed by the binding of ribosome-<u>a</u>ssociated inhibitor A (RaiA), which prevents ribosomal splitting and blocks tRNAs from binding the P- and A-sites of the ribosome (Polikanov *et al*. 2012; Maki and Yoshida 2022).

Structural studies have provided many key insights into dynamics in the ribosome and its interplay with associated RNA and proteins throughout these processes. Often, these approaches involved trapping specific ribosomal states using small molecule inhibitors or genetic mutations (Korostelev 2022), or via biochemical purifications or reconstitutions of desired states (Seffouh *et al*. 2024). Major potential shortcomings of these approaches include capturing off-pathway intermediates, losing weakly associated protein factors during purification, or limiting biologically relevant conformational heterogeneity. In contrast, *in situ* cryogenic electron tomography (cryoET) preserves the native cellular environment, but remains technically challenging, and is both low-throughout and limited in resolution relative to cryoEM-based methods (Dickerson and Lucas 2025). Recognizing these challenges, we and others have developed methods to apply cryoEM to crudly purified cell extracts (Davis *et al*. 2016; Kastritis *et al*. 2017; Verbeke *et al*. 2018; Ho *et al*. 2020), enabling analysis of ribosomes and other macromolecular complexes (Tüting *et al*. 2021; Su *et al*. 2021; Skalidis *et al*. 2022).

Here, we introduce cryoPRISM, a rapid *ex vivo* cryoEM approach combining sample preparation and extensive computational analysis to reconstruct near-atomic resolution cryoEM maps of ribosomes imaged from rapidly clarified *E. coli* cell lysates. Specifically, we establish cell lysis, vitrification, and image analysis methods, using them to visualize 23 unique ribosomal states spanning ribosomal assembly, translation initiation, elongation, quiescence, and a rare intermediate in the trans-translation pathway. Using cryoPRISM, we additionally identify a novel configuration of EF-G on idle ribosomes with hibernation factor RaiA co-bound, and we visualize large subunit precursors simultaneously bound to four unique ribosome assembly factors, previously only observed in an affinity purified sample (Nikolay *et al*. 2021). As our approach preserves interactions with a milieu of endogenous ribosomal factors and detects low-abundance structural intermediates, we anticipate it will be broadly applicable in the discovery of novel ribosomal biology in other cell types and will enable comparisons of ribosomal structural landscapes in cells perturbed pharmacologically or genetically.

## RESULTS

### CryoPRISM rapidly produces high-resolution *ex vivo* structures of ribosomes

In developing cryoPRISM, we aimed to produce high-resolution structures of ribosomes from rapidly clarified lysates (**Figure 1A**), while preserving near-native conditions. To this end, we maintained cryogenic conditions during lysis and minimized sample handling between harvesting the culture and plunge-freezing the lysate (see Methods) on cryoEM grids bearing monolayer graphene as a support surface (Grassetti *et al*. 2023). Application of cryoPRISM to wild-type *E. coli* cells grown at either 37°C or 25°C allowed us to reconstruct density maps of 30S and 50S ribosomal subunits, and 70S ribosomes at high-resolution (2.5Å – 2.6Å GS-FSC), each exhibiting protein and rRNA features consistent with these global resolution estimates (**Figures 1B-D, S1**). Given the high degree of structural heterogeneity we expected to observe in these particles, we analyzed these data using cryoDRGN (Zhong *et al*. 2021; Kinman *et al*. 2023), MAVEn (Sun *et al*. 2023) and SIREn (Kinman *et al*. 2025), aiming to extract both previously reported and novel ribosomal states from the structural ensemble.

**Figure 1.**
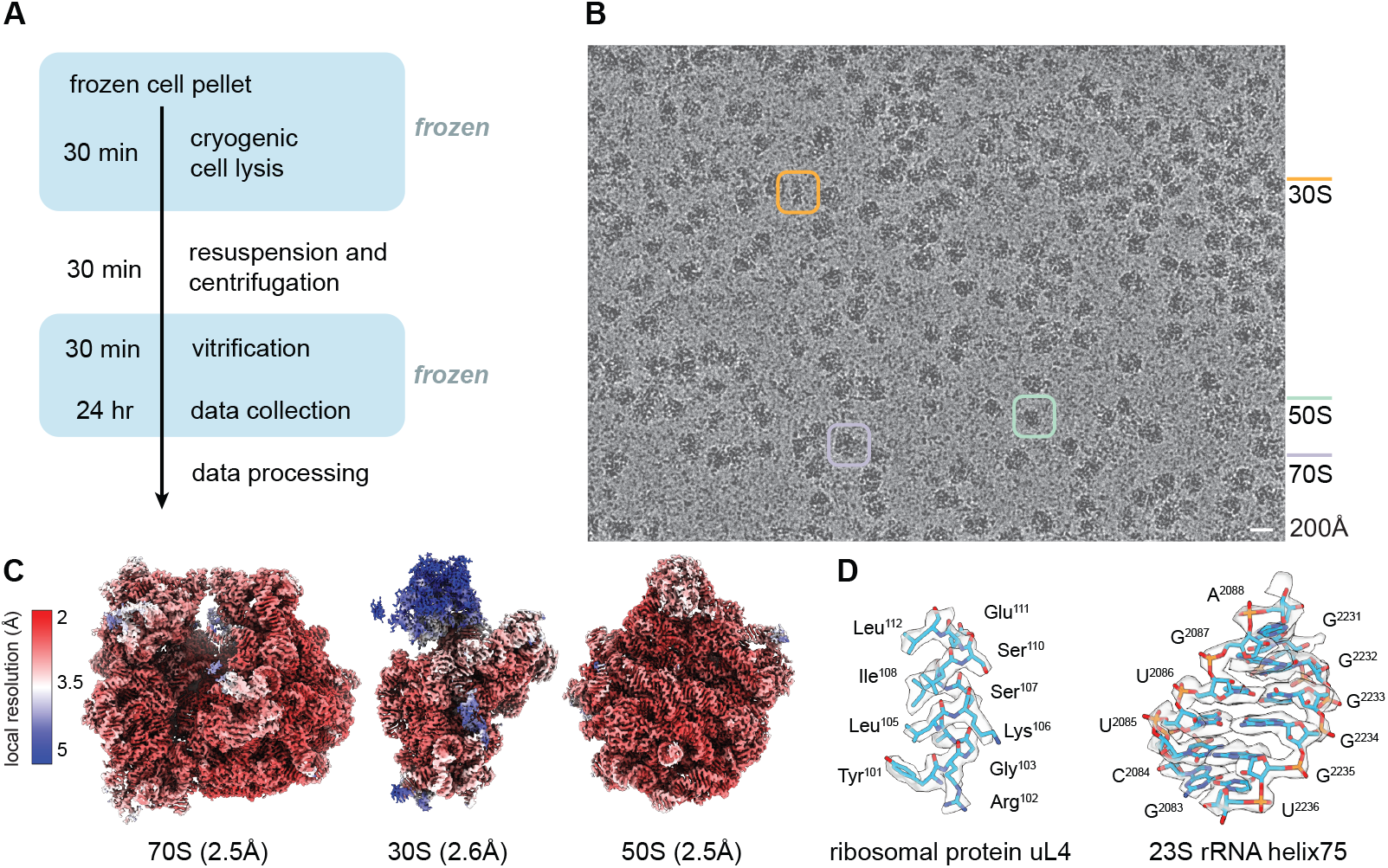
CryoPRISM is a rapid single-particle cryoEM workflow for high-resolution structural analysis of ribosomal particles directly from cell lysates. **(A)** Schematic of the workflow from cell lysis through data collection. Blue boxes indicate periods of time that biological material remains frozen, such that the duration of lysate in solution is minimized. **(B)** Representative micrograph from *E. coli* cells grown at 37°C and prepared following (A). Exemplar 70S, 50S, and 30S particles highlighted in purple, blue, and yellow boxes, respectively. Micrograph is low pass filtered to 5Å; scale bar (white) represents 200Å. **(C)** Reconstructed density maps of 70S, 30S, and 50S particles, colored by local resolution, with scale bar at left. Global resolution estimates based on gold-standard FSC are indicated. **(D)** Density map (grey) reconstructed from 70S particles with the atomic model from PDB 7st6 (Carbone *et al*. 2021) overlaid, highlighting regions of ribosomal protein uL4 and rRNA helix 75, with residues labeled. Atomic model was refined using ISOLDE (Croll *et al*. 2018).

### Translating ribosomes imaged in *E. coli* lysates

We first focused our analysis on 70S particles resolved in cells grown at 37°C, observing actively translating ribosomes, indicated by the presence of mRNA and tRNAs in the classical ‘A/A, P/P, E/E’ and ‘P/P, E/E’ states at frequencies of 20% and 8%, respectively (**Figures 2A, S2A-B**). Although base stacking near the tRNA and mRNA interaction was well resolved, specific mRNA codon-tRNA anticodon pairs were not visualized, likely due to the reconstruction having averaged across an ensemble of distinct mRNA/tRNA pairs. Interestingly, we observed density corresponding to E-site tRNA in all actively translating ribosomes analyzed (**Movie S1**). We could additionally resolve ratcheting of the small subunit relative to the large subunit, consistent with conformational rearrangements coupled to translocation (**Figure 2B**). Hybrid tRNAs in the’A/P*’ and’P/E’ configuration were observed in a minority of particles (∼1%) and only in cells grown at 25°C (**Figure 2A, S2C**).

**Figure 2.**
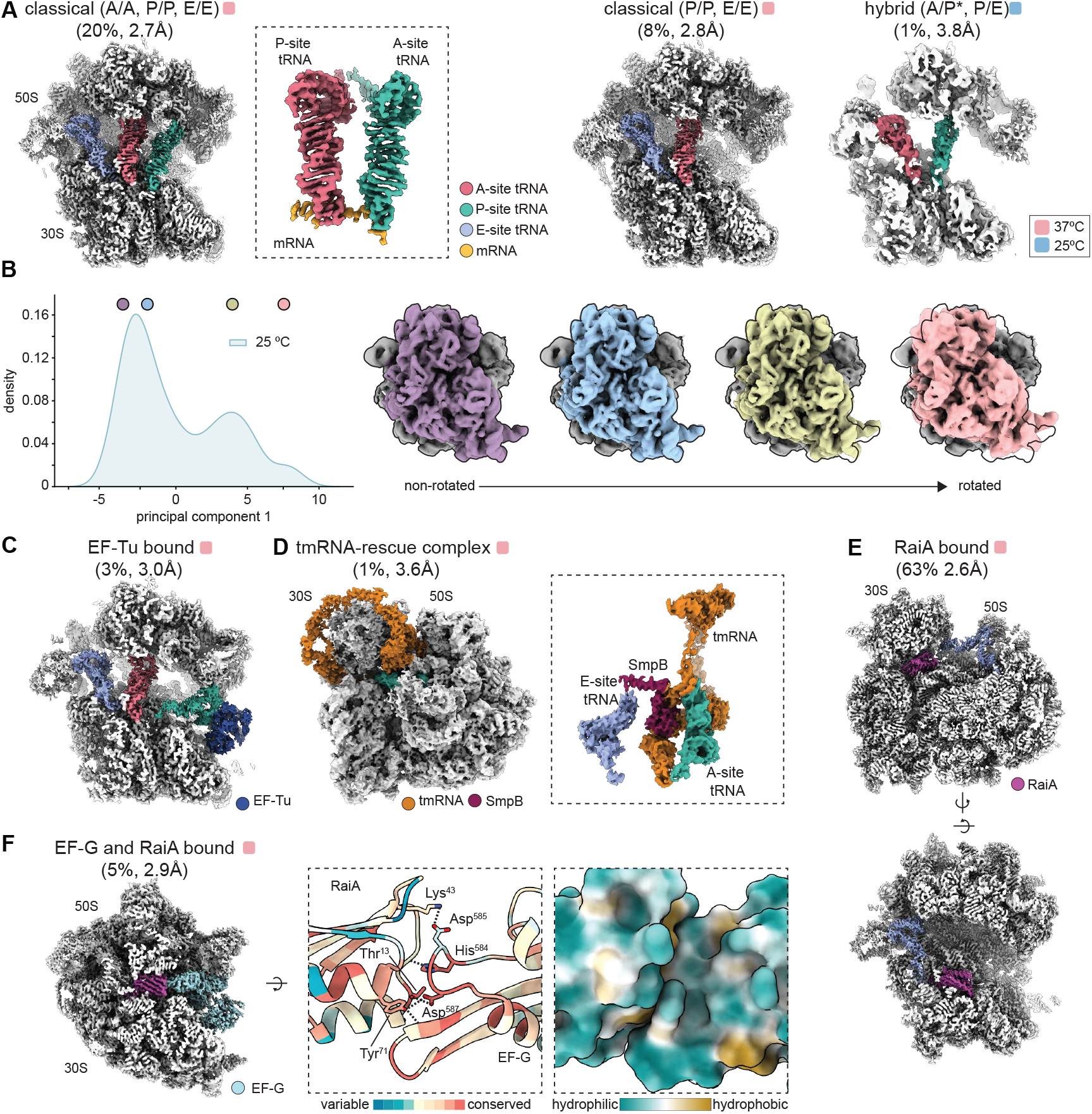
70S ribosomes visualized from *E. coli* cell lysates span diverse functional states involved in translation, rescue, and quiescence. **(A)** Density maps of 70S particles showing ribosomes with tRNAs in the [A/A, P/P, E/E] sites (left) or [P/P, E/E] sites (middle) in the classical conformation, and in the [A/P*, P/E] sites in the hybrid conformation (right). Maps are clipped to show tRNA-binding sites and are colored by chain according to the legend using atomic models 7st6 or 7ssn (Carbone et al. 2021) refined using ISOLDE (Croll 2018). Inset highlights [A/A, P/P] tRNAs resolved in the classical conformation. The percentage of all 70S particles classified into each state and estimated global resolution of each map are noted. The dataset from which each map was resolved is indicated by a pink (37°C) or blue (25°C) square. **(B)** To parameterize conformational changes in 70S particles resolved in the 25°C dataset, 500 cryoDRGN volumes were sampled from *k*-means centroid locations in the latent space. Principal component analysis (PCA) was conducted on voxels within a mask encompassing 16S rRNA helix 6, 10, and 17 (see **Figure S13D**), and the particle density along the first component is shown (left). Density maps were sampled along this axis at locations marked with circles, and the corresponding maps are shown (right), with the large subunit colored gray and the small subunit colored according to the sampled location. An outline of the left-most sampled volume (purple, non-rotated state) is depicted in the same reference as the remaining maps for comparison. **(C)** Density map of a ribosome bound by aa-tRNA·EF-Tu, where EF-Tu is in the compact conformation (dark blue), aa-tRNA (green), P-site tRNA (pink), and E-site tRNA (purple) are colored using PDB 6wd2 (Loveland *et al*. 2020) as in (A). The percentage of 70S particles classified into this state and estimated global resolution of the map are noted. **(D)** Density map of a tmRNA-SmpB complex bound to the ribosome, with SmpB (maroon) and tmRNA (orange) colored using PDB 8vs9 (Teran *et al*. 2024), with additional rigid-body fitting per chain in Phenix. Density observed for A and E-site tRNA colored as in (A). Boxed inset shows an alternative view focused on tRNA binding site. **(E)** Density map of an idle ribosome with RaiA (magenta) bound and E-site tRNA (purple) present, and with A- and P-sites vacant. A clipped alternative view highlights the RaiA binding site. Map colored using 8vty (Aleksandrova et al. 2024) and 7st6, as in (A). **(F)** Density map (left) and built atomic models (right) of EF-G and RaiA simultaneously bound to an idle ribosome. In the density map, RaiA and EF-G are colored magenta and sky blue, respectively. Boxed insets show the RaiA·EF-G interface, with cartoon representation of atomic models colored by evolutionary conservation (left) using ConSurf (Yariv *et al*. 2023), and as a space-filling model colored by hydrophobicity (right) using the ChimeraX implementation of pyMLP (Laguerre *et al*. 1997). Residues within 3Å that appear to interact through hydrogen bonding or salt bridges are labeled, and stick representation colored by atom are shown (left).

We captured a small population (∼3%) of elongating ribosomes during delivery of aminoacyl-tRNA (aa-tRNA) by EF-Tu (**Figure 2C, S2D**), with several features of this intermediate consistent with the initial codon/anticodon discrimination stage of mRNA decoding (Loveland *et al*. 2020). During this stage, amino-acylated tRNA is thought to probe the A-site mRNA for appropriate base pairing prior to conversion of the 30S to a closed conformation and subsequent GTP hydrolysis by EF-Tu (Loveland *et al*. 2020). Consistent with this codon/anticodon discrimination state, we observed: *1)* the 30S subunit in an open conformation relative to the large ribosomal subunit, which placed EF-Tu in its compact state distal to the sarcin-ricin loop (**Figure S3A**); and *2)* ordered switch I and switch II domains in EF-Tu, as expected prior to GTP hydrolysis (**Figure S3B**).

### Direct visualization of a rare intermediate in the trans-translation pathway

In the 37°C dataset, we found that ∼1% of mature ribosomes were bound to tmRNA (**Figure 2D, S2E**), consistent with estimates of trans-translation activity during exponential growth (Moore and Sauer 2005). This structure was resolved at ∼3.6Å resolution and bore clear density for A-site tRNA and tmRNA-SmpB positioned in the P-site, consistent with translation of the first codon on the mRNA-like domain (MLD) of tmRNA having occurred (**Figure 2D, inset**). These ribosomes adopted the non-rotated ratchet state, likely in a state preceding translocation by EF-G, and we observed an extreme tilt (∼19°) in the head domain of the 30S away from the 50S, presumably to accommodate the bulky tmRNA (**Figure S3C**). Interestingly, we again detected E-site tRNA on all tmRNA-bound ribosomes, suggesting that E-site tRNA does not readily dissociate in these conditions.

### Discovery of a novel idle ribosome state bearing EF-G and hibernation factor RaiA

A large population of 70S ribosomes were translationally inactive (*i*.*e*., idle), as indicated by a lack of mRNA and P-site tRNA. Within this population, we resolved two distinct high-resolution structures; one with ribosomes bound only by ribosome-<u>a</u>ssociated inhibitor <u>A</u> (RaiA) and tRNA in the E-site (**Figures 2E, S2F**), and a second that bore additional density around the GTPase associated center (GAC). Using an unbiased structure-to-sequence assignment approach (see Methods), we determined this density corresponded to elongation factor G (EF-G) (**Figure 2F, S2G**). This observation of EF-G bound to an idle ribosome in the presence of RaiA was unexpected. Indeed, whereas EF-G is crucial for translational elongation (Gupta *et al*. 1971) and acts in splitting ribosomes during the ribosomal recycling phase of translational termination (Hirashima and Kaji 1973; Seely and and Gagnon 2022), it was hitherto unknown to associate with idle ribosomes. In fact, RaiA was previously believed to sterically clash with EF-G (Polikanov *et al*. 2012), as highlighted by rigid-body docking of existing EF-G atomic models into our density maps (**Movie S2**).

To understand how these idle ribosomes simultaneously accommodated EF-G and RaiA, we built atomic models of these proteins into our density map (**Figure 2F, Table S1**). Comparing our model to the most similar previously observed conformation revealed profound motions in EF-G domains III, IV, and V relative to domains I and II (**Movie S3**). These motions appear to accommodate RaiA, which was positioned in a previously observed binding site (Polikanov *et al*. 2012; Svetlov *et al*. 2021; Aleksandrova *et al*. 2024). Our structure also revealed a well-packed interface between EF-G and RaiA, supported by a series of hydrogen bonds and a salt bridge between largely conserved residues (**Figure 2F**).

### A temperature-sensitive equilibrium between active and inactive 30S conformations

Using in-cell chemical probing, McGinnis and colleagues previously observed that a substantial fraction of free 30S ribosomal subunits adopt an “inactive” state (McGinnis *et al*. 2015), which led us to ask whether cryoPRISM could resolve such a state in lysates. Briefly, as a key feature of these “inactive” states was an altered conformation of helix 44 (H44), we focused on this region of the structure and observed a mixture of active and inactive particles (**Figure 3A, S4A-C**). Nearly half of the active 30S particles were bound by initiation factor 1 (IF1) and initiation factor 3 (IF3), with IF1 positioned near the A-site, as seen previously (Carter *et al*. 2001), and with IF3 exhibiting its canonical “dumbbell” shape (Hussain *et al*. 2016) with both the N- and C-terminal domains visible. Inactive particles were observed in two states; one with a portion of H44 unlatched and physically separated from the decoding center, and a second with H44 unresolved, presumably a result of excessive flexibility. Strikingly, the distribution of these states was strongly temperature-dependent, with inactive particles making up ∼30% of all small subunit particles at 37°C but decreasing to ∼1% at 25°C (**Figure 3B**). Additionally, at 25°C, a large majority (∼85%) of active particles were bound by IF1 and IF3, whereas this frequency was ∼50% at 37°C (**Figure 3B**). As we did not observe IF1 or IF3 independently bound to 30S particles (**Figure S5**), we concluded that they bind to a mature decoding center cooperatively, as previously reported *in vitro* (Zucker and Hershey 1986).

**Figure 3.**
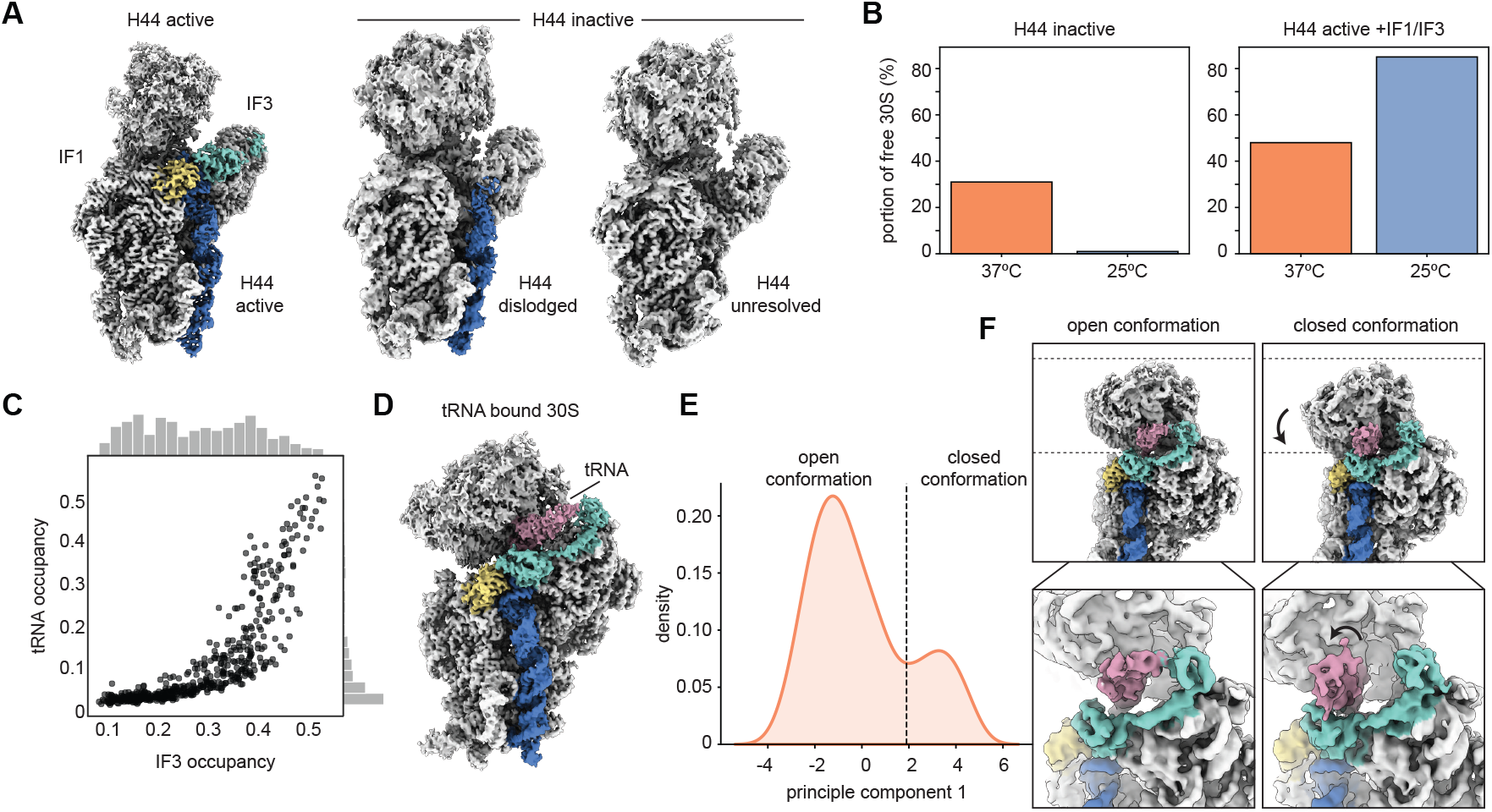
Growth temperature modulates the distribution of active and inactive small ribosomal subunits observed in *E. coli* lysates. **(A)** Density maps of small subunit with H44 (blue) in an active (left), or inactive (middle, right) conformation, from the 37°C dataset. Inactive conformations are distinguished by H44 being dislodged but resolved (middle) or completely unresolved (right). IF1 (yellow) and IF3 (teal) are also resolved in the H44 active density map. Maps are colored using atomic models 7oe1 (Maksimova *et al*. 2021), 8eyq (Sun *et al*. 2023), and 5lmp (Hussain *et al*. 2016), each refined using ISOLDE (Croll 2018). **(B)** Bar chart showing the percentage of H44-inactive small subunit particles in datasets collected from cells grown at 37°C and 25°C (left). H44-inactive particles include both the dislodged and unresolved H44 populations. The percentage of all small subunit particles with H44 in the active conformation and bound by IF1 and IF3 is plotted by cell growth temperature (right). **(C)** Scatter plot depicting the relationship between tRNA and IF3 occupancy, where each dot denotes one of 500 volumes sampled from *k*-means centroid locations in the cryoDRGN latent space. The cryoDRGN model was trained on 30S particles in the 37°C dataset. Histograms illustrate the marginal distributions. **(D)** Density map of the small subunit recovered from the 37°C dataset showing tRNA (purple) in the P-site, as well as IF1, IF3, and H44 clearly resolved and colored as in (A). **(E)** To parameterize conformational changes of P-site tRNA on 30S subunits in the 37°C dataset, 500 cryoDRGN volumes were sampled from the P-site tRNA-positive small subunit particles (see Methods, **Figure S14D**), and principal component analysis (PCA) was conducted on voxels within a mask encompassing the P-site tRNA. The resulting particle density along the first principal component is shown. This component captures a transition of the small subunit with initiator tRNA from an open conformation, with the head tilted away from the body, to a closed conformation, with the head tucked in closer to the body. A threshold used to separate particles for reconstructing the open (PC1 < 1.9) and closed (PC1 ≥ 1.9) conformations, which are shown in (F), is marked with a dotted line. **(F)** Density maps resulting from homogeneous refinements of particles in the open (left) or closed (right) conformation. Maps are low-pass filtered at 5Å. Dotted lines mark common references to better visualize motion. Inset shows a magnified view, focused on the tRNA movement.

### 30S subunits prepare to initiate translation

At 37°C, a subset of active 30S particles contained initiator tRNA in the P-site (**Figure 3C-D, S4D**), with this association dependent on the presence of both IF3 and IF1 (**Figure S5**). Deeper examination using principal component analysis of the volume ensemble (see Methods) revealed a transition from an “open” 30S conformation, where the head was tilted away from the body and the initiator tRNA was unaccommodated, to a “closed” 30S conformation, with the head tucked in closer to the body and the tRNA appearing to transition towards an accommodated state (**Figure 3E-F, Movie S4**). Particles were distributed in a bimodal fashion along this axis of conformational change, consistent with cooperativity in the transition, and high-resolution structures (**Figure S4E-F**) of each could be resolved by pooling particles from the open or closed regions of the distribution (**Figure 3F, top**). The open conformation resembled an *in vitro* reconstituted state proposed to form soon after tRNA binding (Hussain *et al*. 2016) with the P-site tRNA tilted away from the 30S body. In the closed conformation, the P-site tRNA was tilted closer to the body (**Figure 3F, bottom**), likely representing a later intermediate closer to the fully accommodated state (Hussain *et al*. 2016).

### CryoPRISM captures large ribosomal subunit assembly intermediates

We identified an ensemble of large subunit structural states (see Methods), which were distinguished by their overall maturation, rRNA processing status, and the combination of assembly factors bound (**Figures 4A-B, S6-9**). Class A, the most mature, included subclasses either with or without E-site tRNA; Class B exhibited incomplete maturation of the L7/L12 stalk and peptidyl transferase center; and Class C, which lacked density for the central protuberance (CP), captured the least mature particles (**Figure 4A, S6A**). As previously observed in purified large subunits (Davis *et al*. 2016; Sheng *et al*. 2023, Du *et al*. 2024a, Webster *et al*., in preparation), the assembly factor YjgA bound precursors in two conformations: an extended form that occluded the H68 docking site (**Figure 4A, class B2**), and a bent form positioned atop docked H68 (**Figure S9A, class 5**).

**Figure 4.**
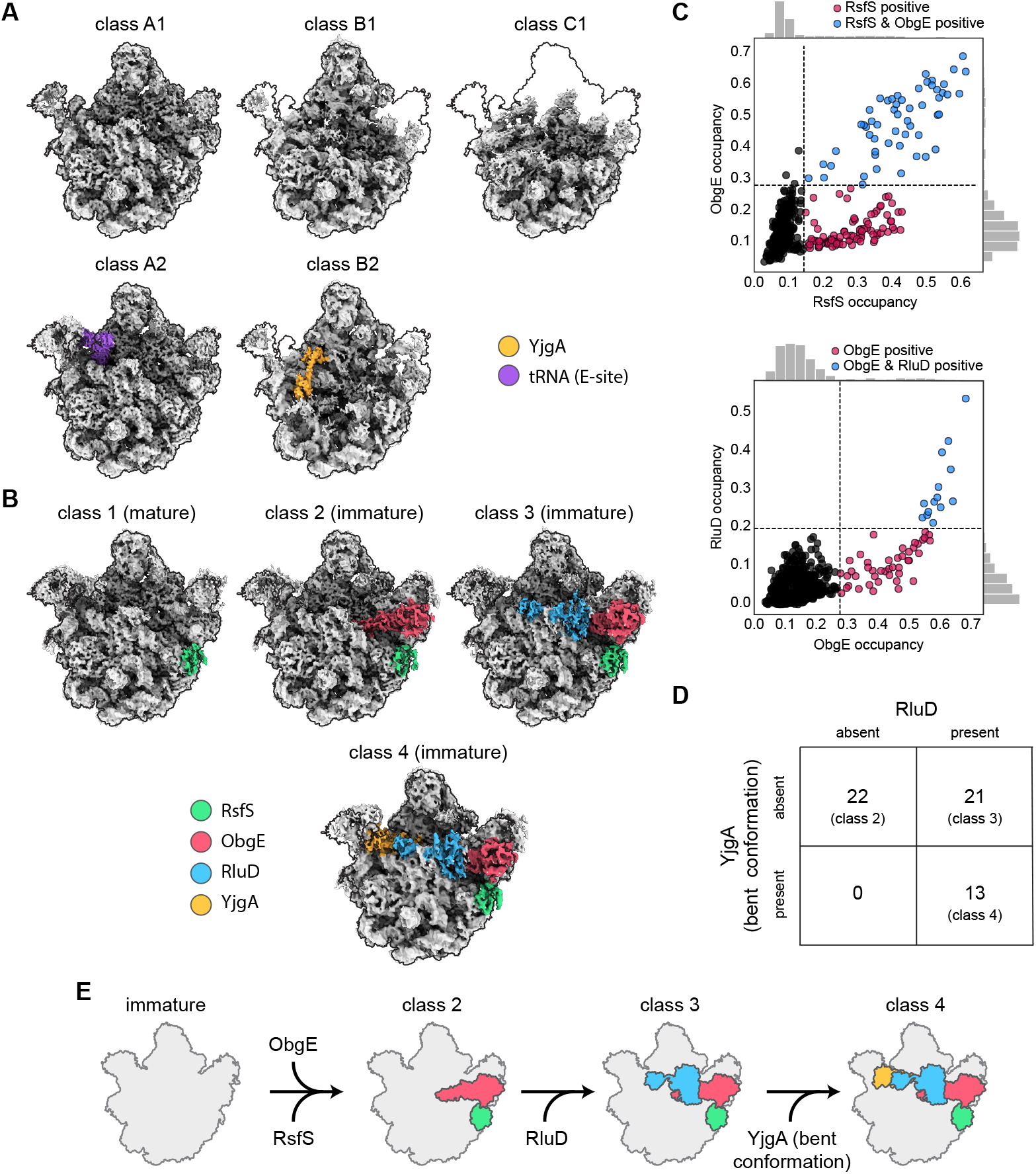
CryoPRISM resolves large subunits assembly intermediates, including those bound to assembly factors in lysates. **(A)** Density maps of major assembly intermediate states resolved in 25°C dataset. For ease of comparison, each density map is outlined using that from class A1. E-site tRNA and YjgA colored purple and yellow, respectively. Maps are colored by atomic models noted in Table S5, each refined with ISOLDE (Croll 2018). **(B)** Density maps of particles bound by the RsfS (green), ObgE (red), RluD (blue), and YjgA (yellow) network of assembly factors. Maps are colored by atomic models noted in Table S4, each refined with ISOLDE (Croll 2018). Reconstructions are outlined as in (A). **(C)** Scatter plots depicting the relationship between RsfS and ObgE (top) and ObgE and RluD (bottom), where each dot denotes one of 500 volumes sampled from *k*-means centroid locations in the cryoDRGN latent space. The cryoDRGN model was trained on 50S particles in the 25°C dataset. Black dots indicate volumes lacking both factors, pink dots indicate volumes with presence of one of the factors, while blue dots indicate volumes with presence of both factors as assessed using expert-defined thresholds noted with dashed lines. Histograms illustrate the marginal distributions. **(D)** Contingency table reflecting presence or absence of YjgA (bent conformation) and RluD in density maps also bearing both RsfS and ObgE. Factor occupancy was assessed via expert-guided inspection of cryoDRGN-generated density maps sampled from *k*-means centroid locations of latent space (*k*=500) from the 25°C dataset. **(E)** Schematic summarizing order of large subunit assembly factor binding, as inferred from occupancy relationships (C-D).

CryoPRISM additionally captured large subunit assembly intermediates simultaneously bound by the assembly factors RsfS, ObgE, RluD, and YjgA (bent conformation). To compare the factor binding dependencies observed in lysates to a previously proposed model of factor binding (Nikolay *et al*. 2021), we leveraged cryoDRGN and MAVEn (Sun *et al*. 2023; Kinman *et al*. 2025) to generate hundreds of density maps and to analyze them for factor co-occupancy patterns (see Methods). ObgE was present only when RsfS was also present, and RluD was observed exclusively in the presence of ObgE (**Figure 4C**). As YjgA binding was dependent on RluD when RsfS and ObgE were bound, we placed it downstream in a sequential pathway for factor binding (**Figure 4D-E**). We also observed an additional population of YjgA-bound precursors lacking RsfS, ObgE, and RluD (**Figure S9A, class 5**), suggesting an independent YjgA entry point. Finally, we observed a novel intermediate in which RsfS and ObgE are bound to particles lacking density for the central protuberance (**Figure S9A, class 6**), arguing that these factors can also act early in the biogenesis pathway, potentially along a parallel branch of assembly (Davis *et al*. 2016).

Many maps contained RsfS alone, which led us to ask whether these particles were mature subunits or assembly intermediates. To distinguish between these possibilities, we assessed the processing status of the 23S rRNA termini in factor-bound states, as fully processed 5’ and 3’ termini are hallmarks of mature particles (Gutgsell and Jain 2012). RsfS-only particles lacked density for unprocessed 23S rRNA ends, consistent with complete rRNA maturation, whereas particles we assigned as assembly intermediates showed clear density for unprocessed nucleotides at both termini (**Figure S9B**). Having assigned states as either assembly intermediates or mature subunits, we next quantified the frequency of assembly-factor bound precursors, finding that growth at low temperature dramatically increased their abundance (**Figure S9C**).

## DISCUSSION

In the past two decades, structural biology has made progress towards visualizing macromolecular complexes in increasingly native contexts (Nogales and Mahamid 2024). CryoPRISM continues this trajectory, aiming to retain the diversity of structural states present in the cell while avoiding the laborious experimental (*i*.*e*., FIB-milling and/ or tilt-series data collection) and computational challenges (*i*.*e*., exhaustive 2D-template matching or tomographic data processing) currently required for *in situ* cryo-EM (Lucas *et al*. 2021; Zheng *et al*. 2024) and cryo-ET (Wan 2025; Carrion and Davis 2025). Using grid preparation and image analysis techniques described here, we captured high-resolution views of more than 20 ribosomal states expected at various stages of the translation cycle, as well as novel structural states in ribosome assembly and hibernation. Taken together, these structures highlighted cryoPRISM’s promise in both visualizing expected states in more native contexts and in facilitating the discovery of new structural states to be subsequently functionally tested.

By leveraging the single-molecule nature of cryo-EM, we were additionally able to quantify the relative abundance of these states, which hints at the relative rates at which these states interconvert in cells or in our lysis conditions, with low abundance or unobserved states likely having rapidly converted to the next stage of translation, and those that accumulated having likely resulted from a slow kinetic transition to the next state. For example, the classical ‘A/A, P/P, E/E’ state was most abundant in our analysis, consistent with prior single-molecule FRET assays reporting that translating ribosomes dwell predominantly in this classical configuration at the relatively high levels of Mg^2+^ used in our buffers (Munro *et al*. 2007; Kim *et al*. 2007; Aitken and Puglisi 2010; Chen *et al*. 2013). Relatedly, we observed an EF-Tu bound elongation intermediate at the initial stage of mRNA decoding, consistent with prior pre-steady-state kinetic studies showing that aa-tRNA·EF-Tu·GTP ternary complex delivery to the ribosome is rapid relative to GTP-hydrolysis induced conformational changes in EF-Tu, particularly when a non-cognate tRNA has been delivered (Pape *et al*. 1998; Pape *et al*. 2000). Notably, we also observed several related pre-accommodation stages during translation initiation, suggesting that slow cognate codon discrimination can drive the accumulation of specific structural states throughout translation.

Comparison between cells grown at different temperatures revealed distributional changes in the abundance of specific states, consistent with a temperature-dependent change in the kinetics of translation elongation. Specifically, the hybrid ‘A/P*, P/E’ state was observed exclusively at 25°C, indicating that at cold-temperatures, its resolution may be slowed or its formation may be expedited relative to that at 37°C. We note that a subset of obligate intermediates along the translation elongation cycle (*e*.*g*., termination complexes) were not observed at either temperature in this study, which could have resulted from any combination of low cellular abundance, instability in our cell conditions, or incompatibility with our vitrification conditions – emphasizing that one may proceed with caution when interpreting the lack of a structural state.

Notably, some low-frequency states could readily be observed with cryoPRISM, including an intermediate in the trans-translation pathway that had been previously resolved via biochemical purification (Fu *et al*. 2010; Mishra *et al*. 2018; Teran *et al*. 2024). Trans-translation involves EF-Tu delivering tmRNA, in complex with SmpB, to the vacant A-site of the stalled ribosome, followed by accommodation of the tRNA-like domain (TLD) in the A-site, where it receives the nascent peptide chain. Subsequently, EF-G catalyzes the translocation of the TLD and SmpB to the P-site and, eventually, the ribosome transitions from the original mRNA template to the mRNA-like domain (MLD) of the tmRNA (Müller *et al*. 2021). In our structure, we observe TLD/SmpB in the P-site with the MLD in the decoding center and an accommodated A-site tRNA. This configuration suggests that translation of the initial codon of the tmRNA reading frame is slow, potentially due to the bulky nature of tmRNA-SmpB that may be challenging to translocate out of the P-site.

Ribosomes bound to RaiA in complex with EF-G were unexpected, particularly at the relatively high (∼5%) abundance observed. However, close inspection revealed a novel, highly ordered interface exhibiting both chemical complementarity and conservation between RaiA and EF-G, consistent with an evolutionarily selected role for this complex. Interestingly, while EF-G has never been observed on hibernating ribosomes in bacteria, EF-Tu has been observed on hibernating ribosomes in *B. subtilis* (Pereira *et al*. 2015) and in the γ-proteobacterium *Psychrobacter urativorans* (Helena-Bueno *et al*. 2024), where the ribosome was also bound to additional hibernation factors. Additionally, EF-G’s eukaryotic analog, eEF2, has a well-established association with translationally inactive ribosomes in both yeast and mammalian cells bound to hibernation factors (Anger *et al*. 2013; Liu and and Qian 2016; Brown *et al*. 2018; Smith *et al*. 2021; Du *et al*. 2024b). In further support of this model of elongation-factor associated hibernating ribosomes, recent *in situ* work in *E. coli* cells found poorly resolved density consistent with a GTPase in the GTPase-associated center of the ribosome on “100S-like” hibernating disomes (Powell *et al*. 2025). Taken together, these observations raise the question of why hibernating ribosomes would associate with elongation factors. While still an area of active research, it has been hypothesized that, by forming complexes with elongation factors, ribosomes may shield their RNase- and protease-sensitive active sites from damage (Helena-Bueno *et al*. 2024), or that the factors themselves are less protease sensitive once bound to a hibernating ribosome (Du *et al*. 2024b). Our observation of bacterial EF-G bound to an idle ribosome contributes to this expanding appreciation of the complexities and regulation of ribosomal hibernation.

Our data also provide a framework for understanding how multiple assembly factors act in a coordinated sequence during late large subunit biogenesis. RsfS is known to associate with large subunits as a silencing factor that prevents subunit joining, acting on either assembly intermediates or mature subunits (Häuser *et al*. 2012; Nikolay *et al*. 2021). ObgE and RluD, in contrast, are implicated in late large subunit assembly; ObgE through structural remodeling driven by GTP hydrolysis and RluD through pseudouridylation of rRNA (Wout *et al*. 2004; Jiang *et al*. 2006; Feng *et al*. 2014; Nikolay *et al*. 2021; Siibak and Remme 2010; Popova and Williamson 2014). Our findings support a model in which RsfS, ObgE, RluD, and YjgA function in concert during the final stages of large subunit maturation. Building on a model proposed by Nikolay and colleagues, our data suggest the following sequence: RsfS and ObgE associate first; RluD joins next; YjgA binds last (**Figure 4E**). Importantly, our structures confirm RsfS can engage both mature and immature particles, consistent with its dual roles in silencing and assembly.

Given the complexity of a cell lysate, we expected to observe other large, abundant macromolecules beyond the ribosome. Consistent with this, we resolved the GroEL chaperonin complex at 3.8Å resolution (**Figure S10**), though other abundant complexes including RNA polymerase and ATP synthase were not observed.

Taken together, we find that cryoPRISM offers a generalizable, high-resolution strategy for interrogating structural landscapes of ribosomes without purification. By preserving endogenous interactions in these clarified lysates and leveraging modern tools to analyze structural heterogeneity in cryoEM datasets, this approach revealed both expected and previously unobserved ribosomal states, ranging from transient decoding intermediates to rare hibernation and assembly complexes. The identification of EF-G on idle ribosomes, the cooperative binding sequence of late-acting assembly factors, and the observation of a particle in the process of trans-translation underscore cryoPRISM’s capacity to capture biologically meaningful transitions. Unlike in situ cryo-ET, cryoPRISM is accessible without needing specialized equipment and is amenable to high-throughput comparative studies, making it especially well-suited for profiling structural dynamics across physiological conditions, developmental stages, and diverse species. As structural biology continues its shift toward increasingly native contexts, cryoPRISM fills a crucial niche by balancing throughput, resolution, and proximity to cellular conditions to discover and quantify structural states of life’s quintessential molecular machine.

## MATERIALS AND METHODS

### Bacterial strains and cell growth

To generate lysates suitable for imaging, *E. coli* strain NCM-3722 (Brown and Jun 2015) with the Marionette array (Meyer *et al*. 2019) integrated into its genome (strain st_ JD886) was grown with aeration at 37°C in LB medium to OD ∼1 and diluted a minimum of 10-fold into 1L fresh LB medium at either 37°C or 25°C and cultured with aeration before harvesting at a cell density of OD600 ∼ 0.5. To harvest, cultures were quenched on ice and cells were pelleted by centrifugation at 4000g for 15 minutes in a Beckman JLA-8.100 rotor. Pellets were scraped in small chunks, flash frozen in liquid nitrogen, and then stored at -80 °C.

### Cell lysis

Frozen cell pellets were lysed under cryogenic conditions using a Retsch MM-400 Mixer Mill with a 5 mL stainless steel grinding jar and 10 mm stainless steel grinding balls, each cooled using liquid nitrogen. Cells were milled for five cycles at 30 Hz for 90 seconds, where the grinding jar was cooled in liquid nitrogen for at least 30 seconds between each cycle. Frozen lysate was then resuspended in ice-cold Buffer A (20mM Tris-HCl, 100mM NH_4_Cl, 10mM MgCl_2_, 0.5mM EDTA, 6mM β-mercaptoethanol, pH 7.5). Lysates were clarified by two sequential centrifugation steps at 4°C: an initial spin at 21,000g for 20 minutes, after which the supernatant was transferred to a new tube with addition of 33 U/mL DNase I and DNase I Reaction Buffer (New England Biolabs #M0303L and #B0303S, respectively), followed by a second spin at 21,000g for 10 minutes. The resulting buffer composition was: 28 mM Tris-HCl, 90mM NH_4_Cl, 11.5mM MgCl_2_, 0.45mM EDTA, 0.5mN CaCl_2_, 5.4mM β-mercaptoethanol, pH 7.5.

### Cryogenic electron microscopy data collection and standard data processing

Following Grassetti and colleagues, 200-mesh Quantifoil R2/1 copper grids were coated with a monolayer of graphene and treated with UV/ozone for 10 minutes using a Bioforce PC440 UV/ozone cleaner (Grassetti *et al*. 2023) followed by immediate application of sample (4μL) to the graphene side of the grid. Grids were blotted using a FEI Vitrobot Mk IV instrument for 4 seconds with a blot force of -2 at 10°C and 95% relative humidity, followed by plunge freezing in liquid ethane. Datasets were collected with EPU aberration-free image shift (AFIS) on a Titan Krios G3i operated at 300kV using a Gatan K3 detector equipped with a BioQuantum energy filter. Imaging parameters included: a total dose of 45.2 e-/Å fractionated across 45 frames; defocus range from -0.03 to -2 μm in steps of -0.2 μm; and magnification of 81,000x. Movies (9,748 for 37°C dataset, and 7,908 for the 25°C dataset) were collected in super-resolution mode and binned 2X, resulting in a nominal pixel size of 1.06Å.

Data pre-processing, including motion correction and contrast transfer function estimation, particle picking, and particle extraction (box size of 440 pixels) were performed in Warp 1.0.9 (Tegunov and Cramer 2019), producing 1,503,396 and 1,003,018 particles for the 37°C and 25°C datasets, respectively. Particle stacks were processed in cryoSPARC v4.5.3 (Punjani *et al*. 2017) following depicted scheme (**Figures S11-S12**), and using default parameters for listed jobs, unless otherwise specified. Briefly, particles were subjected to *heterogenous refinement* using six input volumes: a mature ribosome, small subunit, and large subunit, and three “junk” volumes that were hand-picked from a 6-class *ab-initio* job run on all imported particles. Particle stacks that sorted into the mature ribosome, small subunit, or large subunit classes from the heterogenous refinement job were then subjected to several rounds of additional filtering via multiclass *ab-initio*, where particles sorting into “junk” *ab-initio* output classes were excluded with each round. This process resulted in initial particle stacks for mature ribosomes (37°C:516,905 particles; 25°C:261,520 particles), small subunits (37°C:155,227; 25°C:125,742), and large subunits (37°C:271,184; 25°C:132,075).

### CryoDRGN training

The initial particle stacks for each ribosomal class where further filtered and analyzed for heterogeneity using cryoDRGN v2.2 (Zhong *et al*. 2021). Briefly, particle poses and estimated CTF parameters were extracted from refinements described above, and particles were downsampled to box size of 128 pixels (3.64Å/pixel) using cryoDRGN pose and ctf parsing, and downsampling commands. CryoDRGN models were trained using the following hyperparameters: encoder and decoder network architectures:256×3 nodes; learning rate:1e-4; minibatch size:8 epochs:25. The resulting cryoDRGN model was used to filter ‘junk’ particles by removing *k*-means clusters visually determined to be poorly posed or undesired particles (Kinman *et al*. 2023). Filtered particle stacks were used for *ab-initio reconstruction* and *homogenous refinement* in cryoSPARC, then subjected to a second round of cryoDRGN model training, using the same parameters as describe above (**Figures S11-S12**).

### Occupancy analyses

#### MAVEn ‘on-the-fly’ analysis

Volumes were generated by cryoDRGN using latent embeddings for each particle at box size 64 and occupancy within masks corresponding to the feature of interest were queried for each generated volume. Queried maps were binarized using SIREn’s CNN-predicted binarization thresholds using the default model, as described previously (Kinman *et al*. 2025). In all instances where masks were derived from atomic models, they were generated using the `molmap` command in ChimeraX (Meng *et al*. 2023) and converted to a mask using the `relion_mask_create` command to binarize and apply a soft-edge using default parameters unless otherwise specified. Final occupancy values were computed by summing voxels within each masked region in the queried volume relative to the mask itself, resulting in a fractional occupancy for each structural feature analyzed. Thresholds to distinguish particles with low-occupancy versus high-occupancy for a structural feature were defined based on manual inspection of representative volumes above or below that cutoff.

#### MAVEn ‘k-means’ analysis

Analyses were performed as above with the following changes: *1)* volumes were generated at each *k*-means centroid location (as opposed to for every particle); *2)* volumes were generated at box size 128; *3)* when performing refinements using subsets identified by the occupancy analysis, all particles assigned to a class in the selected set were pooled.

### 70S analysis

The 37°C and 25°C 70S datasets were filtered by magnitude of the latent variable to remove remaining particles that produced poor reconstructions, reducing the 37°C particle stack from 455,260 to 451,636 particles and 25°C from 252,575 to 247,684, respectively, and producing highly featured latent spaces for each dataset (**Figure S13A**). In this space for the 37°C dataset, tmRNA-bound particles formed a distinct cluster (**Figure S13A**) and were directly selected and used for *ab-initio reconstruction* followed by *homogeneous refinement* in cryoSPARC (**Figure 2D**).

The EF-Tu bound state (**Figure 2C**) and the EF-G-RaiA-bound state (**Figure 2F**) were analyzed as depicted (**Figure S13B**). First, these particles were identified using MAVEn *‘on-the-fly’* (Sun *et al*. 2023) as described above using masks corresponding to a factor in the GTPase-associated center (GAC), and to the P/P-site tRNA (**Figure S13B**). Particles with a fractional occupancy greater than 0.2 for a GAC-bound factor and 0.35 for P/P-site tRNA were considered positive for these factors, whereas particles below these thresholds were considered negative. This resulted in 20,147 particles with both a GAC-bound factor and P/P-site tRNA, and 39,563 particles with only a GAC-bound factor and no P/P-site tRNA. Both particle stacks were exported to cryoSPARC and *ab-initio reconstructions* were used as input models for *homogeneous refinements*. Subsequent 3D-classification with a mask around the GAC was used to further separate particles based on subtle differences that could not be detected at box size 64 – specifically discriminating between EF-G and EF-Tu bound ribosomes. Among the P/P-site tRNA positive particles, two out of three classes corresponded to EF-Tu and were pooled for a final homogeneous refinement in cryoSPARC. The third class exhibited GTPase density more closely resembling EF-G but was not further refined due to low confidence. Among the P/P-site tRNA absent particles, 3D-classification separated a class of EF-G bound ribosomes from a second class exhibiting low-resolution density that was excluded from further refinement due to low resolution. Notably, the EF-G bound particles lacking P-site tRNA contained clear density for RaiA (**Figure 2F**).

After excluding tmRNA or GTPase-bound particles, the remaining particle stack was used to generate 1,000 volumes sampled from *k*-means cluster centers from the latent space and analyzed using MAVEn (Sun *et al*. 2023) at box 128. Masks for A/A, P/P, and E/E-site tRNA were used to separate the classical ‘A/A, P/P, E/E’ and ‘P/P, E/E’ populations (**Figure 2A**), following scheme in Figure S13C. The remaining particles contained high E/E-site tRNA and RaiA occupancy, and they were refined via *homogeneous refinement* in cryoSPARC (**Figure 2E**).

The hybrid tRNA state (**Figure 2A**) was obtained from the 25°C dataset using the full 70S particle stack (252,575 particles; **Figure S12**) and sampling 200 volumes from the *k*-means centroid locations of the latent embeddings. Volumes with hybrid tRNA density were manually selected and corresponding particles (4,830 particles) were used as input for *ab-initio reconstruction* and *homogenous refinement* in cryoSPARC. Subsequent 3D-classification using an A/P*-site tRNA mask separated particles lacking A/P*-tRNA from those with both A/P* and P/E-site tRNAs; the latter was used for *ab-initio reconstruction* and *homogenous refinement* in cryoSPARC (**Figure S13E**).

### 30S analysis

Particles stacks from the 37°C and 25°C datasets corresponding to free 30S subunits were again filtered by magnitude of the latent variable, reducing the particle stack from 133,631 to 119,389 particles and 116,295 to 100,164, respectively (**Figure S14A-B**). To identify the presence, absence, or conformational changes of structural features, including H44, IF1, IF3, and P-site tRNA, we used MAVEn ‘*on-the-fly*’ occupancy analysis, as described above. Briefly, different conformations of H44 on the 16S rRNA were classified using masks specifically designed to capture the difference between an active and a dislodged H44 (**Figure S14C**). Thresholds for designating different 30S states are listed in **Table S2** and **Table S3** for 37°C and 25°C datasets, respectively.

Volumes reconstructed from the 37°C dataset with a fractional occupancy greater than 0.4 for P-site tRNA were used to designate an initial particle stack for tRNA-bound 30S subunits consisting of 23,969 particles (**Figure S14D**). These particles were used for *ab-initio reconstruction* and *homogenous refinement* in cryoSPARC, followed by training of a cryoDRGN model using the same parameters listed above. This particle stack was further filtered using both magnitude of the latent variable and visual quality of volumes sampled from *k*-means centroid locations in latent space, reducing it to 19,195 particles (**Figure S14D**). 500 volumes were then sampled from *k*-means centroid locations of the latent embeddings and analyzed using the described MAVEn ‘*k-means’ analysis*, using a mask encompassing different tRNA conformations (**Figure S14E**). Volumes with a fractional occupancy greater than 0.27, corresponding to 18,090 particles, were designated to have significant tRNA presence. These particles were used for *ab-initio reconstruction* and *homogenous refinement* in cryoSAPRC (**Figure 3D**). To resolve different tRNA configurations, vPCA was performed using the same tRNA mask (**Figure S14E**). To separate particles based on the bimodal distribution along the first principal component, particles below a threshold of 1.9 were used for refinement of the open conformation 30S, and particles above a threshold of 1.9 were used for refinement of the closed conformation of 30S particles (**Figure 3F**).

### 50S analysis

The 50S particle stack from the 25°C dataset (118,843 particles, **Figure S12**) was filtered using volumes sampled from *k*-means centroid locations to remove clusters of poorly resolved particles, resulting in a refined dataset of 114,750 particles (**Figure S15A**). MAVEn *‘k-means’ analysis* was applied to this refined particle stack to determine occupancy dependency relationships between ribosome assembly factors, RsfS, ObgE, RluD, and YjgA (bent conformation) (**Figure 4C-D**). Expert-guided curation was used to classify the presence of YjgA (bent conformation) and RluD for each volume containing RsfS and ObgE (**Figure 4D**). To isolate particle stacks for reconstructing factor-bound states (**Figure 4B, S9A**), MAVEn *‘on-the-fly’ analysis* was applied using masks corresponding to r-protein uL5, rRNA helix H68, E-site t-RNA, RsfS, ObgE, RluD and YjgA (bent conformation), defined as above (**Figure S15B-C**). Thresholds defining each class and PDBs used for each mask are listed in **Table S4**. To examine the structural landscape of the other 50S assembly intermediates, MAVEn *‘k-means’ analysis* was applied to the remaining 93,921 particles, visualized in a heatmap (**Figure S6A**). To isolate particle stacks for reconstructing classes A1, A2, B1, B2, and C1 (**Figure 4A**), MAVEn *‘on-the-fly’ analysis* was applied using masks corresponding to r-proteins uL5 and uL16, rRNA helices H68 and H89, as well as YjgA (extended conformation), RsfS, and E-site tRNA (**Figure S15D-E**). Thresholds defining each class and PDBs used for each mask are listed in **Table S5**. Note that for all reconstructions, particles used in one state were excluded from subsequent selection rounds to ensure all refined particles stacks were strictly non-overlapping.

### Unbiased structure-to-sequence assignment of EF-G

Density in the GTPase region of the ribosome was identified as EF-G based on *de novo* models built by ModelAngelo v1.0 (Jamali *et al*. 2024) using the build_ no_seq function, where chains were fed into a hidden Markov model and assigned to the *E. coli* (K12) proteome (Uniprot ID UP00625) using probability-based scoring.

### Model building

The EF-G·RaiA model was built using Coot (version 0.9.8.96), ChimeraX (version 1.9), and Phenix (version 1.21.2-5419), starting with an AlphaFold3 (Abramson *et al*. 2024) model of *E. coli* RaiA and EF-G. EF-G from this model was then segmented per-domain (residues 1-293; domain I; 294-412: domain II; 413-488: domain III; 489-609: domain IV; 610-700: domain V). Each domain was docked into our EF-G:RaiA density map as a rigid-body, followed by iterative rounds of refinement using phenix.refine and Coot (Casanal *et al*. 2020). Conservation was predicted using the ConSurf web server (Yariv *et al*. 2023). Surface coloring based on hydrophobicity was performed using the ChimeraX implementation of pyMLP (Laguerre *et al*. 1997).

## Supporting information

Movie S1. Actively translating ribosomes contain E/E-site tRNAs.

Movie S2. Existing atomic models of 70S-bound EF-G are incompatible with RaiA binding.

Movie S3. Large domain-wise rearrangements in EF-G facilitate binding to RaiA in idle 70S particles.

Movie S4. Visualization of small subunit transition from open to closed conformation, as captured by principal component analysis of an ensemble of de

## DATA AVAILABILITY

Upon publication, the raw electron micrographs will be publicly deposited at EMPIAR. Density maps and atomic models described in Table S1 will be available through EMDB and the PDB, and cryoDRGN model weights and analysis files will be available through Zenodo.

## AUTHOR CONTRIBUTIONS

Conceptualization: M.M., J.H.D.

Investigation: M.M., G.S.L.P.

Writing–original draft: M.M., G.S.L.P.

Writing–review and editing: all authors

Visualization: all authors

Supervision: J.H.D.

Project administration and funding acquisition: J.H.D.

## ACKNOWLEDGEMENTS

We thank A. Grassetti and B. Sauer for support with graphene-coated grid fabrication and model building, respectively. We thank S. Sterling and J. Podgorski for assistance with cryoEM sample preparation and data collection, and L. Kinman for her feedback throughout the completion of this work. This research was supported by the National Institutes of Health grant R01-GM144542 (JHD), the National Science Foundation GRFP (MM) and CAREER grant 2046778 (JHD), and the Smith Family Odyssey Award (JHD). Samples were prepared at the Automated Cryogenic Electron Microscopy Facility in MIT.nano and screened on a Talos Arctica microscope, which was a gift from the Arnold and Mabel Beckman Foundation.

