## Supplementary Figures and Tables for "Capturing ribosomal structures in cellular extracts with cryoPRISM: A purification-free cryoEM approach reveals novel structural states"

1. Department of Biology
2. Computational and Systems Biology Graduate Program

Massachusetts Institute of Technology,
Cambridge, Massachusetts, 02139, United States.

* equal contributions

### SUPPLEMENTARY MOVIES

**Movie S1. Actively translating ribosomes contain E/E-site tRNAs.**
Using a cryoDRGN model trained on all actively translating ribosome from the 37°C dataset (**Figures 2A,C**) twenty cryoDRGN maps were sampled from *k*-means centroid locations in the cryoDRGN latent space. A/A, P/P, and E/E-site tRNAs are indicated.

**Movie S2. Existing atomic models of 70S-bound EF-G are incompatible with RaiA binding.**
Our atomic model (9pyc) of EF-G (light blue) and RaiA (magenta) overlaid with previously deposited *E. coli* EF-G structures (7ug7, 7pjy, 7pjv, 7n2c, 4v9p,4v9o, 4v7d, 4v7b, 3ja1, 3j9z, 7pjz, 2rdo, 7pjw, 1zn0, 7k54, 1jqm, 3j0e, 4v6t; various colors). Each overlaid EF-G model was docked as a rigid body by fitting the atomic model for domains I and II (residues 1-412) to our density map of idle ribosomes bound by both RaiA and EF-G.

**Movie S3. Large domain-wise rearrangements in EF-G facilitate binding to RaiA in 70S idle ribosomes.**
Conformational changes between EF-G from 7pjz (Petrychenko *et al.* 2021) and our model (9pyc) involves re-arrangement of domains III (residues 413-488), IV (residues 489-609), and V (residues 610-700) by ~7Å, ~12Å, and ~14Å, respectively. Atomic model 7pjz was selected as a starting model due to overall structural similarity to our built model, calculated by RMSD of common residues 100-700 after aligning on domains I and II (residues 1-412). Conformational changes between the two PDBs are visualized using the ChimeraX "morph" feature.

**Movie S4. Visualization of the small subunit transition from an open to closed conformation, as captured by principal component analysis of an ensemble of density maps.** Twenty-five cryoDRGN volumes sampled along the first principal component, as analyzed in Figure 3E. Briefly, 500 cryoDRGN volumes were sampled from the latent space of the P-site tRNA positive 30S particles, and principal component analysis (PCA) was then conducted on voxels within a mask encompassing P-site tRNA. The cryoDRGN volumes shown were sampled by following the observed distribution along the first principal component.

### SUPPLEMENTARY TABLES

| Model and data deposition information | |
|---|---|
| **Model name** | **EF-G·RaiA** |
| **PDB ID** | 9pyc |
| **EMDB ID** | 72030 |
| **EMPIAR ID** | to be deposited |
| **Data collection and image processing** | |
| **Microscope** | Titan Krios G3i |
| **Camera** | Gatan K3 (counting mode) |
| **Magnification (nominal)** | 81,000 X |
| **Accelerating voltage (keV)** | 300 |
| **Total electron dose ($e^-$/Å$^2$)** | 45.18 |
| **Defocus range (µm)** | -0.2 to -2; -0.03 |
| **Micrographs collected** | 9,748 |
| **Pixel size (Å)** | 1.06 |
| **Map reconstruction** | |
| **Image processing package** | cryoSPARC |
| **Final particle count** | 21,018 |
| **Symmetry imposed** | C1 |
| **Resolution (Å)** | |
| **0.143 GSFSC unmasked** | 3.9 |
| **0.143 GSFSC tight mask** | 2.9 |
| **Model composition** | |
| **Non-hydrogen atoms** | 5,679 |
| **Protein residues** | 732 |
| **Ligands** | 0 |
| **Model refinement** | |
| **Refinement package** | Phenix and Coot |
| **Map-to-model cross correlation** | |
| **masked (region built)** | 0.73 |
| **unmasked (entire map)** | 0.17 |
| **RMSD bond lengths (Å) [#>4σ]** | 0.001 [0] |
| **RMSD bond angles (°) [#>4σ]** | 0.404 [0] |
| **Model validation** | |
| **MolProbity score** | 1.10 |
| **Clash score** | 1.49 |
| **C-beta outliers (%)** | n/a |
| **Rotamer outliers (%)** | 0.0 |
| **Ramachandran favored (%)** | 96.7 |
| **Q-Score [mean]** | |
| **RaiA** | 0.53 |
| **EF-G** | 0.37 |

**Table S1.** Cryo-EM data collection, processing, model building, and validation statistics.

| **state** | **ribosomal rRNA** | | **initiation factors** | |
|---|---|---|---|---|
| | H44in (*) | H44out (*) | IF1 (5lmp) | IF3 (5lmp) |
| **H44 active** | ≥ 0.8 | | ≥ 0.475 | ≥ 0.4 |
| | > 0.6 and < 0.8 | ≤ 0.575 | ≥ 0.475 | ≥ 0.4 |
| **H44 dislodged** | > 0.6 and < 0.8 | > 0.575 | | |
| **H44 unresolved** | ≤ 0.6 | | | |

**Table S2.** Occupancy thresholds used in MAVEn '*on-the-fly*' analysis to define populations that make each 30S state from the 37°C dataset, with resulting reconstructions depicted in Figures 3A and S14. Masks for each element (columns) were derived from listed atomic models. Masks used for H44in and H44out, noted with *, are depicted in Figure S14. Note that the H44 active conformation was defined by the union of the conditions listed in rows 1-2.

| **state** | **ribosomal rRNA** | | **initiation factors** | |
|---|---|---|---|---|
| | H44in (*) | H44out (*) | IF1 (5lmp) | IF3 (5lmp) |
| **H44 active** | ≥ 0.8 | | ≥ 0.3 | ≥ 0.25 |
| | | ≤ 0.65 | ≥ 0.3 | ≥ 0.25 |
| **H44 dislodged** | < 0.8 | > 0.65 | | |

**Table S3.** Occupancy thresholds used in MAVEn '*on-the-fly*' analysis to define populations that make each 30S state from the 25°C dataset. Masks for each element (columns) were derived from listed atomic models. Masks used for H44in and H44out, noted with *, are depicted in Figure S14. Note that the H44 active conformation was defined by the union of the conditions listed in rows 1-2.

| class | assembly factors | | | | rRNA | r-protein | tRNA |
|---|---|---|---|---|---|---|---|
| | RsfS (7bl5) | ObgE (7bl5) | RluD (7bl5) | YjgA (7bl5) | H68 (4ybb) | uL5 (4ybb) | E-site (7st6) |
| **1 (RsfS+)** | ≥ 0.45 | ≤ 0.25 | | | ≥ 0.5 | ≥ 0.4 | |
| **2 (RsfS+ ObgE+)** | ≥ 0.35 | ≥ 0.45 | < 0.16 | | | ≥ 0.4 | |
| **3 (RsfS+ ObgE+ RluD+)** | ≥ 0.2 | ≥ 0.3 | ≥ 0.3 | ≤ 0.6 | | | |
| **4 (RsfS+ ObgE+ RluD+ YjgA+)** | ≥ 0.2 | ≥ 0.4 | ≥ 0.2 | ≥ 0.4 | ≥ 0.6 | | ≤ 0.65 |
| **5 (YjgA+)** | | ≤ 0.4 | | ≥ 0.4 | ≥ 0.6 | ≥ 0.4 | ≤ 0.6 |
| **6 (RsfS+ ObgE+ CP-)** | ≥ 0.2 | ≥ 0.3 | | | | < 0.7 | |

**Table S4.** Occupancy thresholds used in MAVEn 'on-the-fly' analysis to define populations in each assembly-factor bound class, with resulting reconstructions depicted in Figure S15. Masks for each element (columns) were derived from listed atomic models.

| class | assembly factor | | rRNA | | r-protein | | tRNA |
|---|---|---|---|---|---|---|---|
| | RsfS (7bl5) | YjgA(2p0t) | H68 (4ybb) | H89 (4ybb) | uL5 (4ybb) | uL16 (4ybb) | E-site (7st6) |
| **A1** | ≤ 0.3 | | ≥ 0.5 | | | ≥ 0.5 | ≤ 0.4 |
| **A2** | ≤ 0.3 | | ≥ 0.5 | | | ≥ 0.5 | ≥ 0.6 |
| **B1** | ≤ 0.3 | | ≤ 0.3 | ≤ 0.35 | | ≤ 0.3 | |
| **B2** | ≤ 0.3 | ≥ 0.65 | ≤ 0.5 | | | | |
| **C1** | | | | | ≤ 0.3 | | ≤ 0.1 |

**Table S5.** Occupancy thresholds used in MAVEn 'on-the-fly' analysis to define populations in each assembly intermediate class, with resulting reconstructions shown in Figure S15.

### SUPPLEMENTARY FIGURES

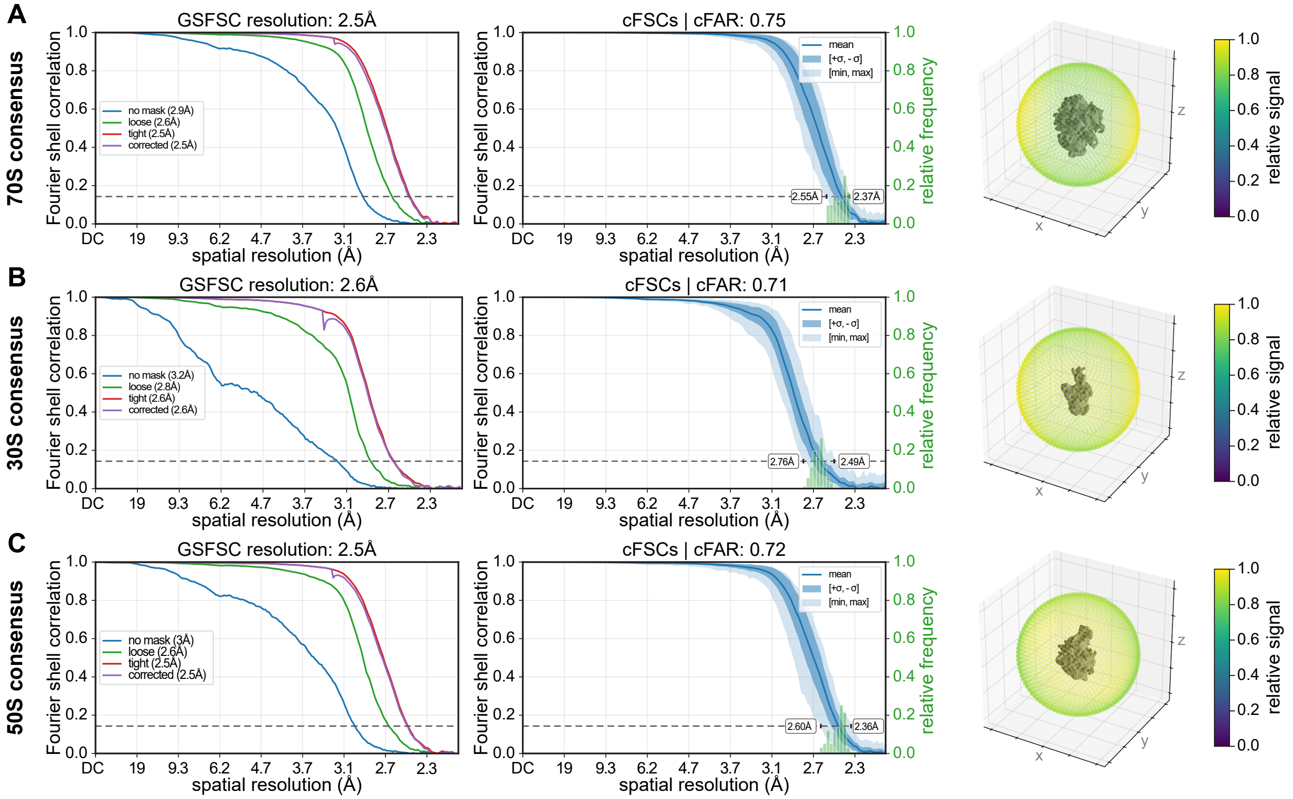


**Figure S1. Quality assessment of consensus 3D reconstructions of 70S, 30S, and 50S ribosomal particles.**
Global resolution estimates calculated using Fourier shell correlation (FSC) between independently refined half-maps as implemented in cryoSPARC's non-uniform refinement method. The FSC = 0.143 threshold is indicated with a dashed line, and the corresponding resolution is reported (left). Conical FSC (cFSC) curves (mean, min, max, ±1σ, as listed in legend) are plotted, with the cFSC area ratio (cFAR) shown in the title and the directional resolution range labeled on the figure. The relative frequency of cFSC values across individual conical slices is shown as a histogram (middle). A sphere colored by relative signal is visualized surrounding each density map (right). All metrics were computed in cryoSPARC from the non-uniform consensus refinements depicted in Figure 1. Plots are shown for: **(A)** the 70S ribosome; **(B)** the 30S subunit; and **(C)** the 50S subunit.

---

**Figure S2. Quality assessment of consensus 3D reconstructions of 70S states (*figure on following page*).**
Global resolution estimates calculated using Fourier shell correlation (FSC) between independently refined half-maps as implemented in cryoSPARC's homogeneous refinement method. The FSC = 0.143 threshold is indicated with a dashed line, and the corresponding resolution is reported (left). Conical FSC (cFSC) curves (mean, min, max, ±1σ, as listed in legend) are plotted, with the cFSC area ratio (cFAR) shown in the title and the directional resolution range labeled on the figure. The relative frequency of cFSC values across individual conical slices is shown as a histogram (middle). A sphere colored by relative signal is visualized surrounding each density map (right). All metrics were computed in cryoSPARC from the homogeneous refinements depicted in Figure 2. Plots are shown for **(A)** the classical (A/A, P/P, E/E) state; **(B)** the classical (P/P, E/E) state; **(C)** the hybrid (A/P*, P/E) state; **(D)** the EF-Tu bound state; **(E)** the tmRNA-rescue complex; **(F)** the RaiA bound state; and **(G)** the EF-G and RaiA bound state.

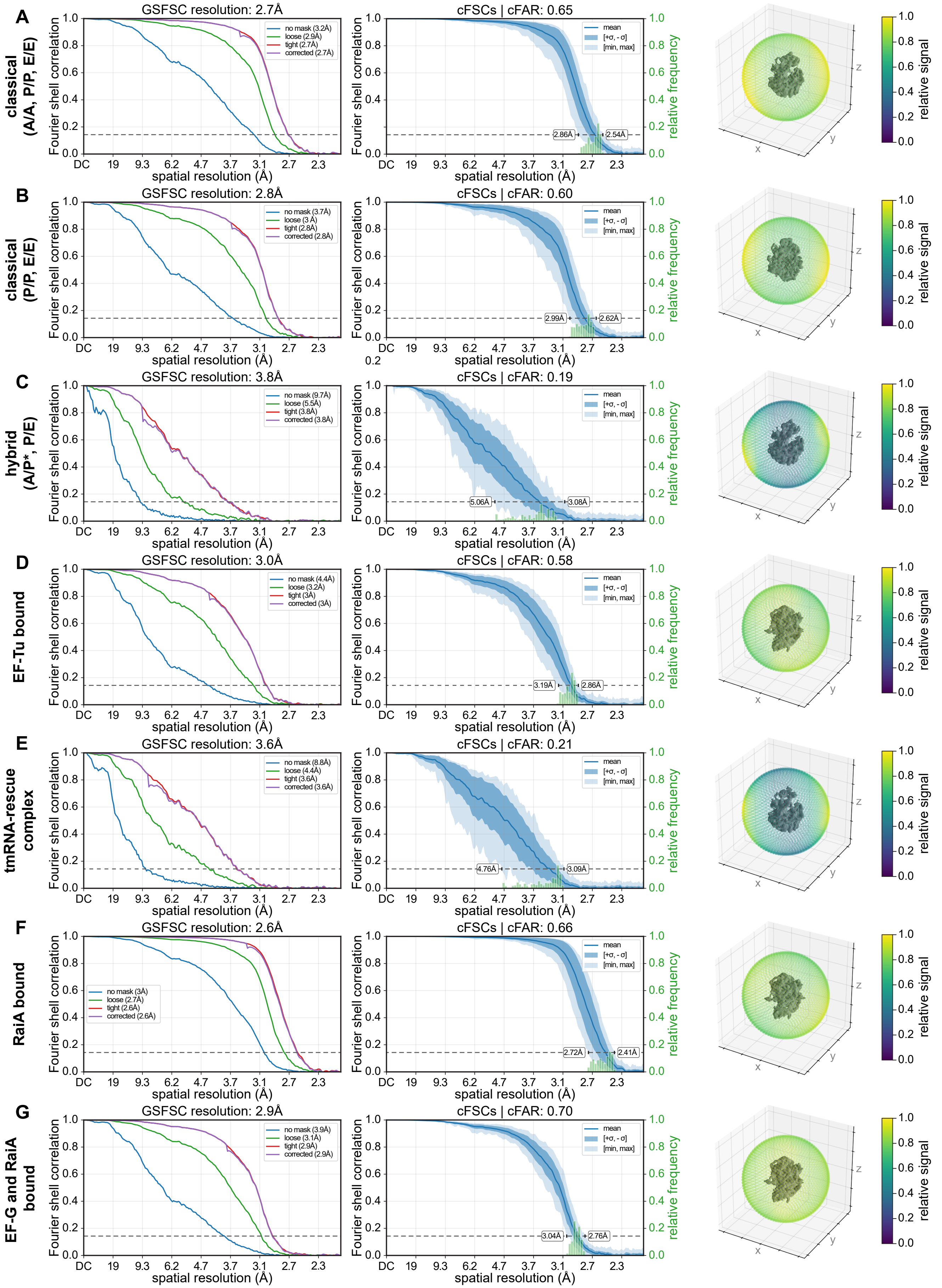

A
classical (A/A, P/P, E/E)
GSFSC resolution: 2.7Å
no mask (3.2Å)
loose (2.9Å)
tight (2.7Å)
corrected (2.7Å)
cFSCs | cFAR: 0.65
2.86Å
2.54Å
B
classical (P/P, E/E)
GSFSC resolution: 2.8Å
no mask (3.7Å)
loose (3 Å)
tight (2.8Å)
corrected (2.8Å)
cFSCs | cFAR: 0.60
2.99Å
2.62Å
C
hybrid (A/P*, P/E)
GSFSC resolution: 3.8Å
no mask (9.7Å)
loose (5.5Å)
tight (3.8Å)
corrected (3.8Å)
cFSCs | cFAR: 0.19
5.06Å
3.08Å
D
EF-Tu bound
GSFSC resolution: 3.0Å
no mask (4.4Å)
loose (3.2Å)
tight (3Å)
corrected (3Å)
cFSCs | cFAR: 0.58
3.19Å
2.86Å
E
tmRNA-rescue complex
GSFSC resolution: 3.6Å
no mask (8.8Å)
loose (4.4Å)
tight (3.6Å)
corrected (3.6Å)
cFSCs | cFAR: 0.21
4.76Å
3.09Å
F
RaiA bound
GSFSC resolution: 2.6Å
no mask (3Å)
loose (2.7Å)
tight (2.6Å)
corrected (2.6Å)
cFSCs | cFAR: 0.66
2.72Å
2.41Å
G
EF-G and RaiA bound
GSFSC resolution: 2.9Å
no mask (3.9Å)
loose (3.1Å)
tight (2.9Å)
corrected (2.9Å)
cFSCs | cFAR: 0.70
3.04Å
2.76Å
Fourier shell correlation
spatial resolution (Å)
DC 19 9.3 6.2 4.7 3.7 3.1 2.7 2.3
mean
[+σ, - σ]
[min, max]
relative frequency
relative signal
x
y
z

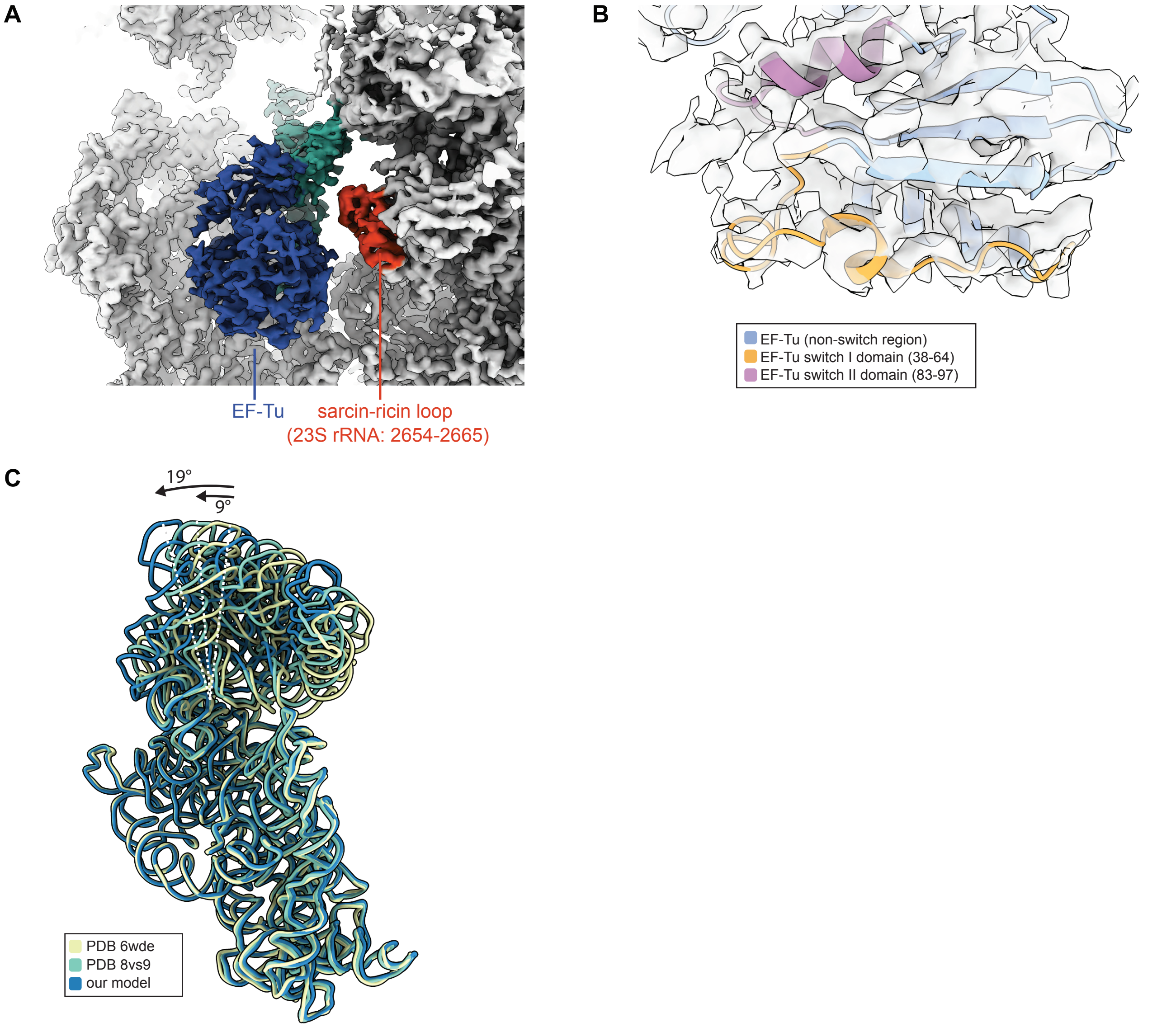


**Figure S3. Structural details of EF-Tu bound 70S ribosome and tmRNA-rescue complex.**

**(A)** Density map of EF-Tu, colored in dark blue, relative to the sarcin-ricin loop (23S rRNA nucleotides 2654-2665), colored in orange, with tRNA in green. Map is colored following atomic model 6wd2 (Loveland *et al.* 2020), as in Figure 2C, and is clipped to better expose EF-Tu.

**(B)** Zoned density map (translucent grey) showing EF-Tu overlaid with atomic model 6wd2, rendered as a cartoon. EF-Tu exhibits ordered switch I (residues 38-64, orange) and switch II (residues 83-97, purple) regions, consistent with a GTP-bound state.

**(C)** Ribbon representations of atomic models 6wde (Loveland *et al.* 2020), 8vs9 (Teran *et al.* 2024), and model generated by segmenting atomic model 8vs9 by rRNA domain and rigid-body docking each domain into our tmRNA-bound map. Ribbons are colored as shown in figure legend. Arrows and dashed white lines indicate the tilt angle of the small subunit head relative to the canonical elongation complex in 6wde (Loveland *et al.* 2020). The angle was approximated using residue 1133 (helix 39) as a reference for head displacement and helix 28 as the pivot point.

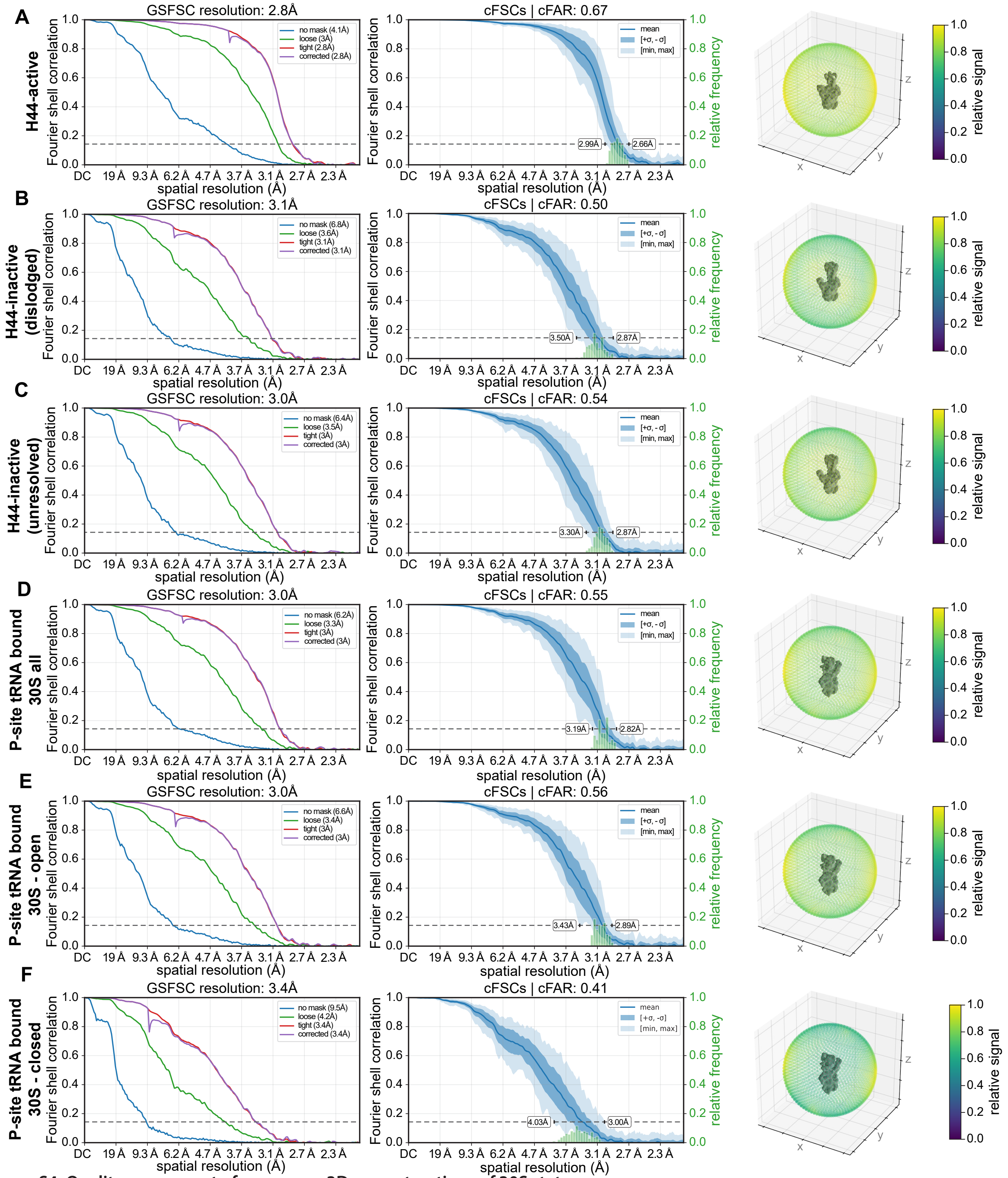


**Figure S4. Quality assessment of consensus 3D reconstructions of 30S states.**

Global resolution estimates calculated using Fourier shell correlation (FSC) between independently refined half-maps as implemented in cryoSPARC's homogeneous refinement method. The FSC = 0.143 threshold is indicated with a dashed line, and the corresponding resolution is reported (left). Conical FSC (cFSC) curves (mean, min, max, ±1σ, as listed in legend) are plotted, with the cFSC area ratio (cFAR) shown in the title and the directional resolution range labeled on the figure. The relative frequency of cFSC values across individual conical slices is shown as a histogram (middle). A sphere colored by relative signal is visualized surrounding each density map (right). All metrics were computed in cryoSPARC from the homogeneous refinements depicted in Figure 3. Plots are shown for **(A)** the H44-active state; **(B)** the H44-inactive dislodged state; **(C)** H44-inactive unresolved state; **(D)** P-site tRNA bound state; **(E)** P-site tRNA bound in an open conformation substate **(F)** P-site tRNA bound in a closed conformation substate.

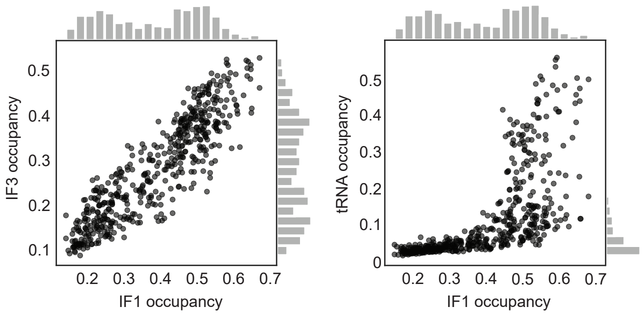


**Figure S5. Relationship between tRNA and initiation factor binding to 30S particles.**
Scatter plot depicting the relationship between occupancy of IF3 and IF1 (left), and P-site tRNA and IF1 (right), where each dot denotes one of 500 volumes sampled from *k*-means centroid locations in the cryoDRGN latent space. The cryoDRGN model was trained on 30S particles in the 37°C dataset. Histograms illustrate the marginal distributions.

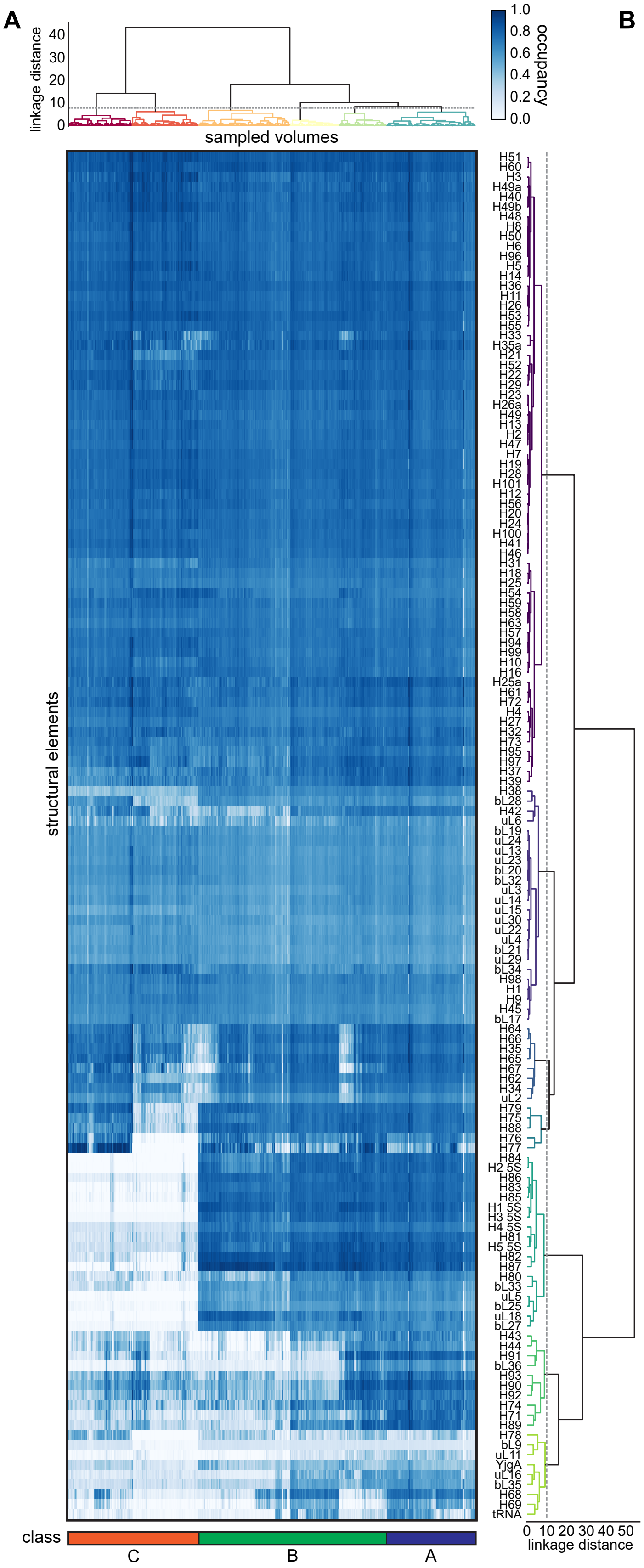


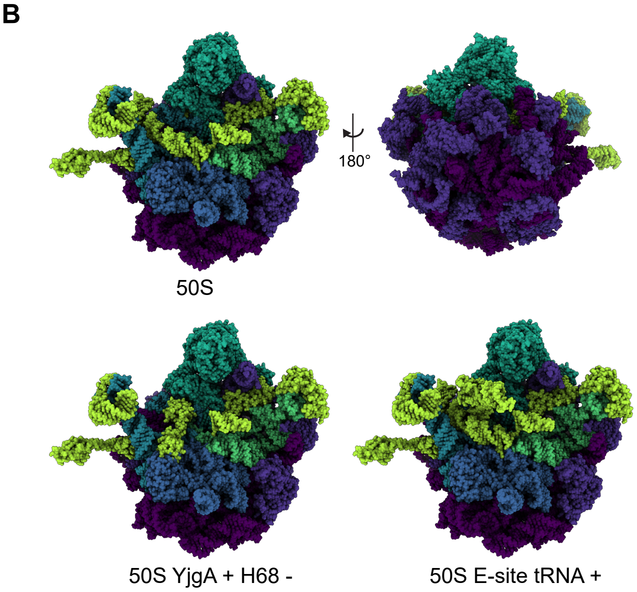


**Figure S6. MAVEn analysis of large subunit particles reveals three major assembly classes.**

**(A)** MAVEn analysis (Sun *et al*. 2023) of large subunit ribosomes displayed as heatmap. Columns represent 500 *k*-means center volumes, and rows represent 138 structural elements derived from atomic models: mature large subunit (4ybb; Noeske *et al.* 2015), YjgA extended conformation (2p0t; Davis *et al.* 2016) and E-site tRNA (7st6; Carbone *et al.* 2021). Hierarchical clustering with the indicated threshold (grey dashed line) partitioned volumes into three classes labeled A-C from most to least mature.

**(B)** Atomic models used in the MAVEn analysis, colored by structural blocks defined by hierarchical clustering in (A). Structures include the mature large ribosomal subunit (4ybb), the large subunit with YjgA (extended conformation) and helix H68 missing (4ybb modified to include 2p0t), and the 50S subunit with E-site tRNA (7st6).

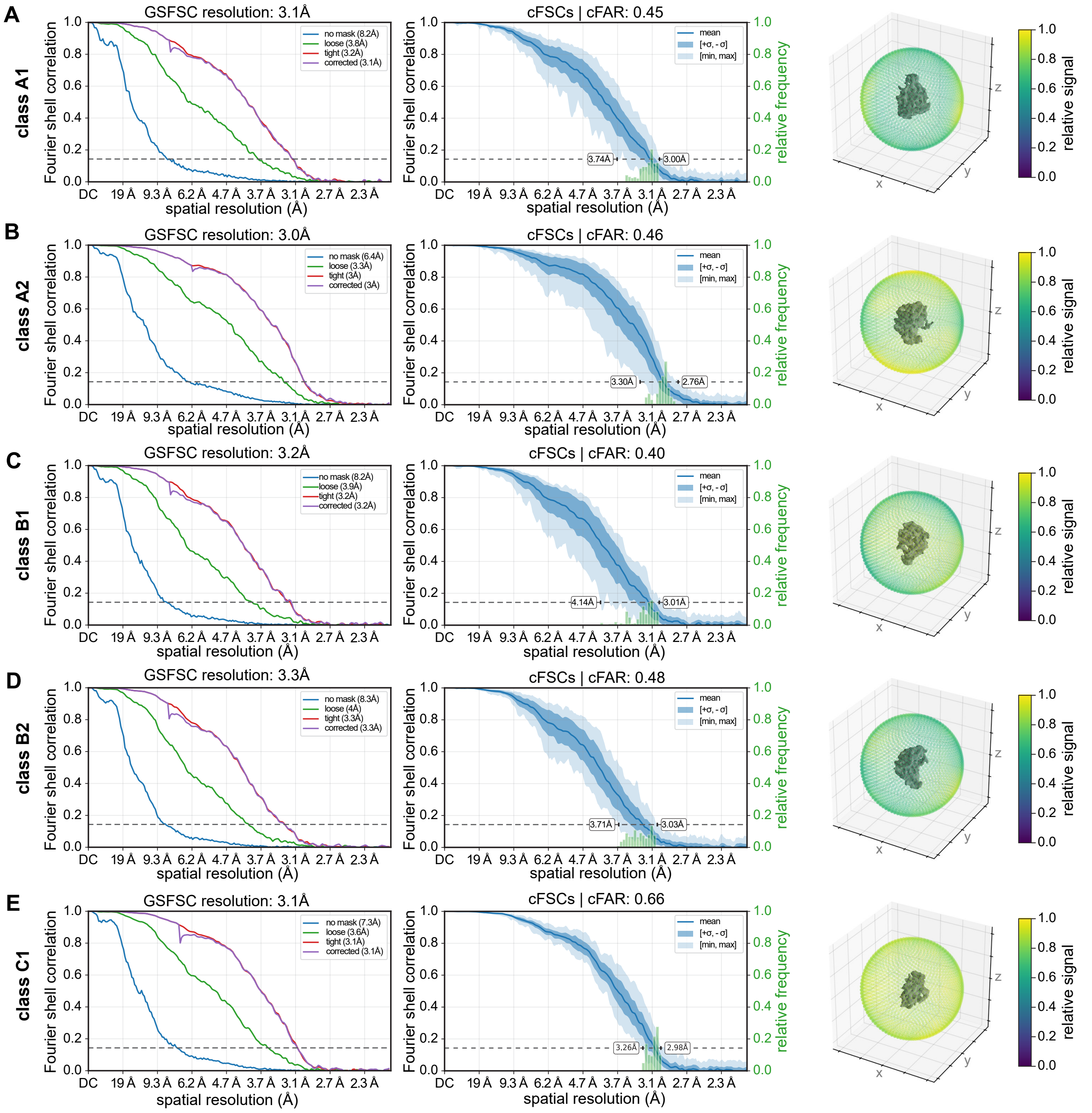


**Figure S7. Quality assessment of consensus 3D reconstructions of large subunit assembly intermediates A1-C1.**
Global resolution estimates calculated using Fourier shell correlation (FSC) between independently refined half-maps as implemented in cryoSPARC's homogeneous refinement method. The FSC = 0.143 threshold is indicated with a dashed line, and the corresponding resolution is reported (left). Conical FSC (cFSC) curves (mean, min, max, ±1σ, as listed in legend) are plotted, with the cFSC area ratio (cFAR) shown in the title and the directional resolution range labeled on the figure. The relative frequency of cFSC values across individual conical slices is shown as a histogram (middle). A sphere colored by relative signal is visualized surrounding each density map (right). All metrics were computed in cryoSPARC from the homogeneous refinements depicted in Figure 4. Plots are shown for **(A)** class A1; **(B)** class A2; **(C)** class B1; **(D)** class B2; **(E)** class C1.

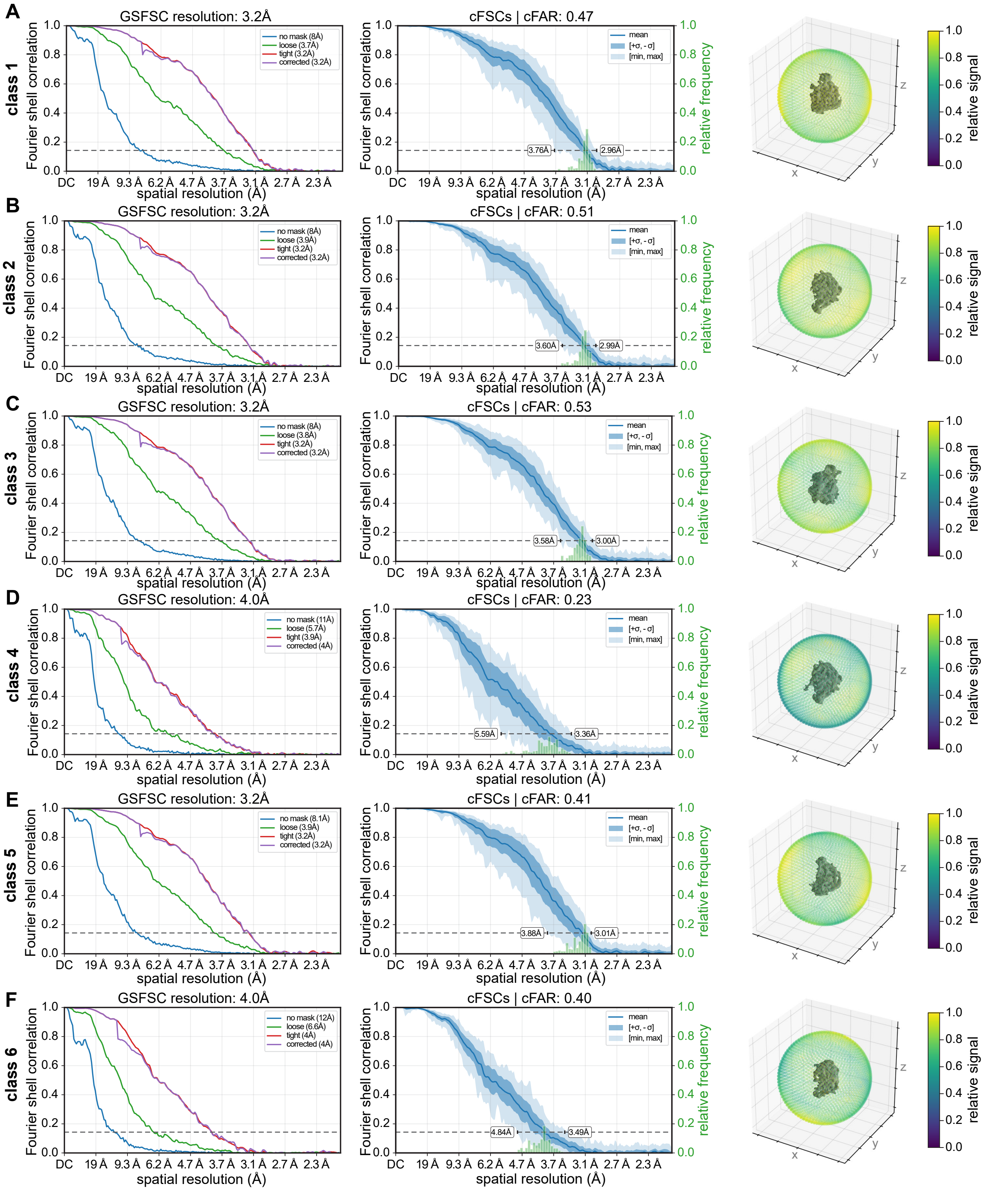


**Figure S8. Quality assessment of consensus 3D reconstructions of large subunit factor-bound classes 1-6.**
Global resolution estimates calculated using Fourier shell correlation (FSC) between independently refined half-maps as implemented in cryoSPARC's homogeneous refinement method. The FSC = 0.143 threshold is indicated with a dashed line, and the corresponding resolution is reported (left). Conical FSC (cFSC) curves (mean, min, max, ±1σ, as listed in legend) are plotted, with the cFSC area ratio (cFAR) shown in the title and the directional resolution range labeled on the figure. The relative frequency of cFSC values across individual conical slices is shown as a histogram (middle). A sphere colored by relative signal is visualized surrounding each density map (right). All metrics were computed in cryoSPARC from the homogeneous refinements depicted in Figures 4 and S9A. Plots are shown for **(A)** class 1; **(B)** class 2; **(C)** class 3; **(D)** class 4; **(E)** class 5; and **(F)** class 6.

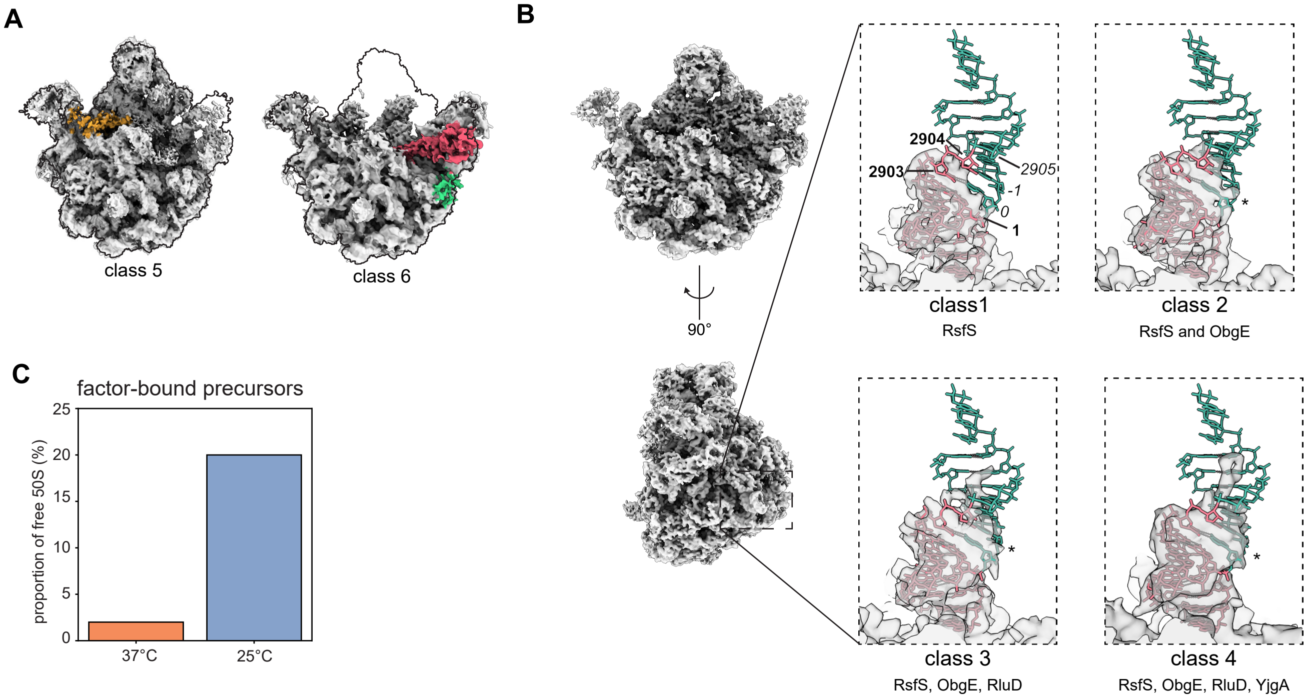


**Figure S9. Detailed structural analysis of factor-bound large subunit particles.**

**(A)** Density maps bearing YjgA (yellow) in a bent conformation (left; class 5), or RsfS (green) and ObgE (red) bound to a large subunit lacking the central protuberance (right; class 6). For ease of comparison, density maps are outlined following the perimeter of class A1 (Figure 4A).

**(B)** Density maps (translucent gray) of classes 1-4 overlaid with stick representation of atomic model 7bl2 (Nikolay et al. 2021) highlighting the 5' and 3' termini of the 23S rRNA. Nucleotides are colored pink for mature, fully processed rRNA termini and green for unprocessed segments, with labels in bold and italics for mature and unprocessed termini, respectively. Asterisk indicates density corresponding to unprocessed termini in classes 2 – 4.

**(C)** Bar chart showing the percentage of 50S particles with ribosome assembly factors bound in the 37°C versus 25°C dataset, as defined by MAVEn '*on-the-fly*' occupancy analysis using masks for RsfS, ObgE, RluD, YjgA (extended conformation), and YjgA (bent conformation). Particles bound only by RsfS were excluded, as these were determined to be mature ribosomal subunits based on the analysis of rRNA termini processing status detailed in panel B.

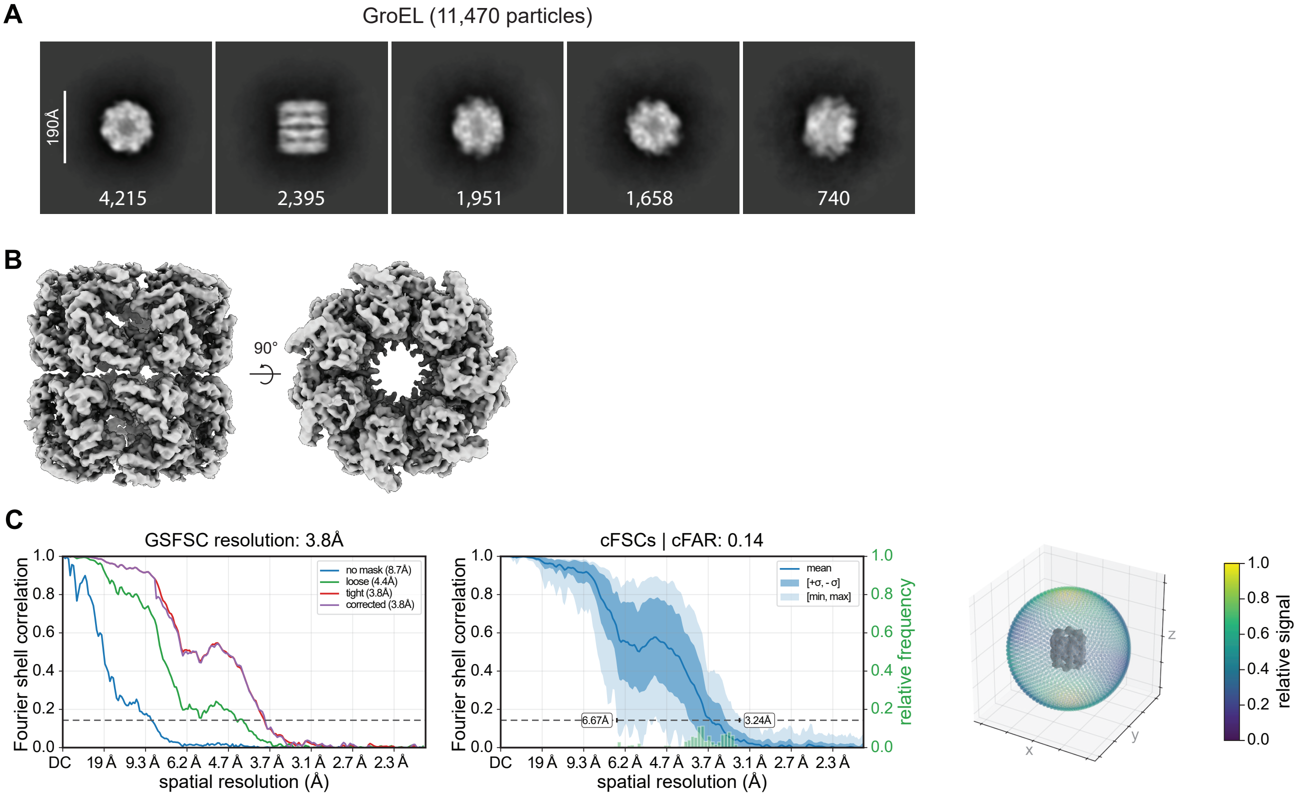


**Figure S10. GroEL resolved using cryoPRISM.**

Particles that sorted into "junk" classes during heterogenous refinement of all extracted particles from the 37°C dataset (Figure S11) were further analyzed using iterative 2D classification, multi-class ab-initio reconstruction, and subsequent homogeneous refinement without symmetry, revealing GroEL.

**(A)** 2D-classes bearing GroEL particles resolved following workflow above. Particle number in each class listed, with a remaining 551 particles partitioning amongst low-resolution classes not shown.

**(B)** Reconstructed density map of GroEL shown from side and top views.

**(C)** Estimates of global resolution, directional resolution, and relative signal in GroEL reconstruction. Masked global resolution estimates were calculated using Fourier shell correlation (FSC) between independently refined half-maps as implemented in cryoSPARC's homogeneous refinement method. The FSC = 0.143 threshold is indicated with a dashed line, and the corresponding resolution is reported (left). Conical FSC (cFSC) curves (mean, min, max, ±1σ, as listed in legend) are plotted, with the cFSC area ratio (cFAR) shown in the title and the directional resolution range labeled on the figure. The relative frequency of cFSC values across individual conical slices is shown as a histogram (middle). A sphere colored by relative signal is visualized surrounding the GroEL density map (right). Note that preferred orientation, which manifested as an anisotropic map, was observed.

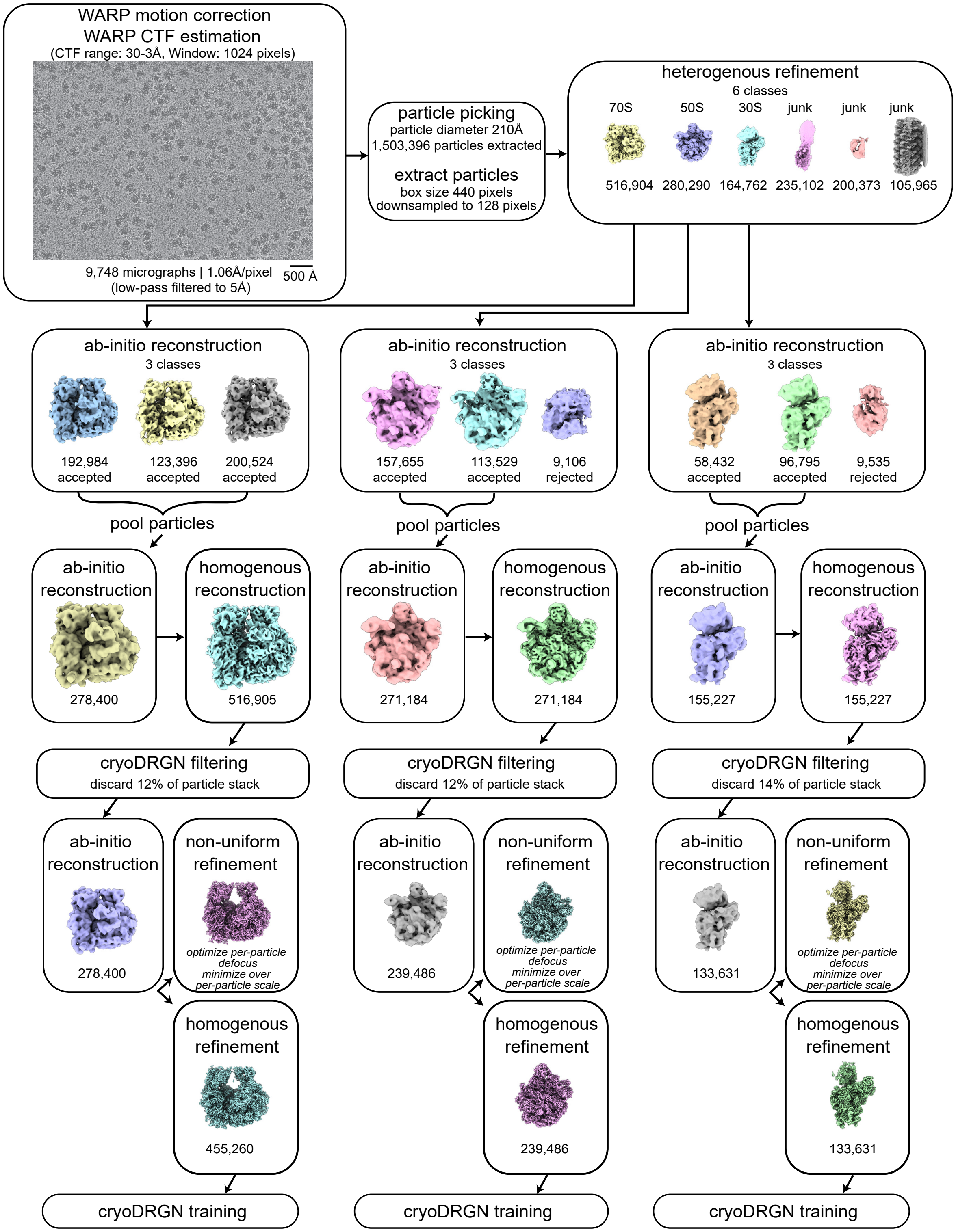


**Figure S11. CryoSPARC pre-processing workflow for analysis of the 37°C dataset.**
The job names and key parameters are listed, with particle numbers listed and representative maps depicted.

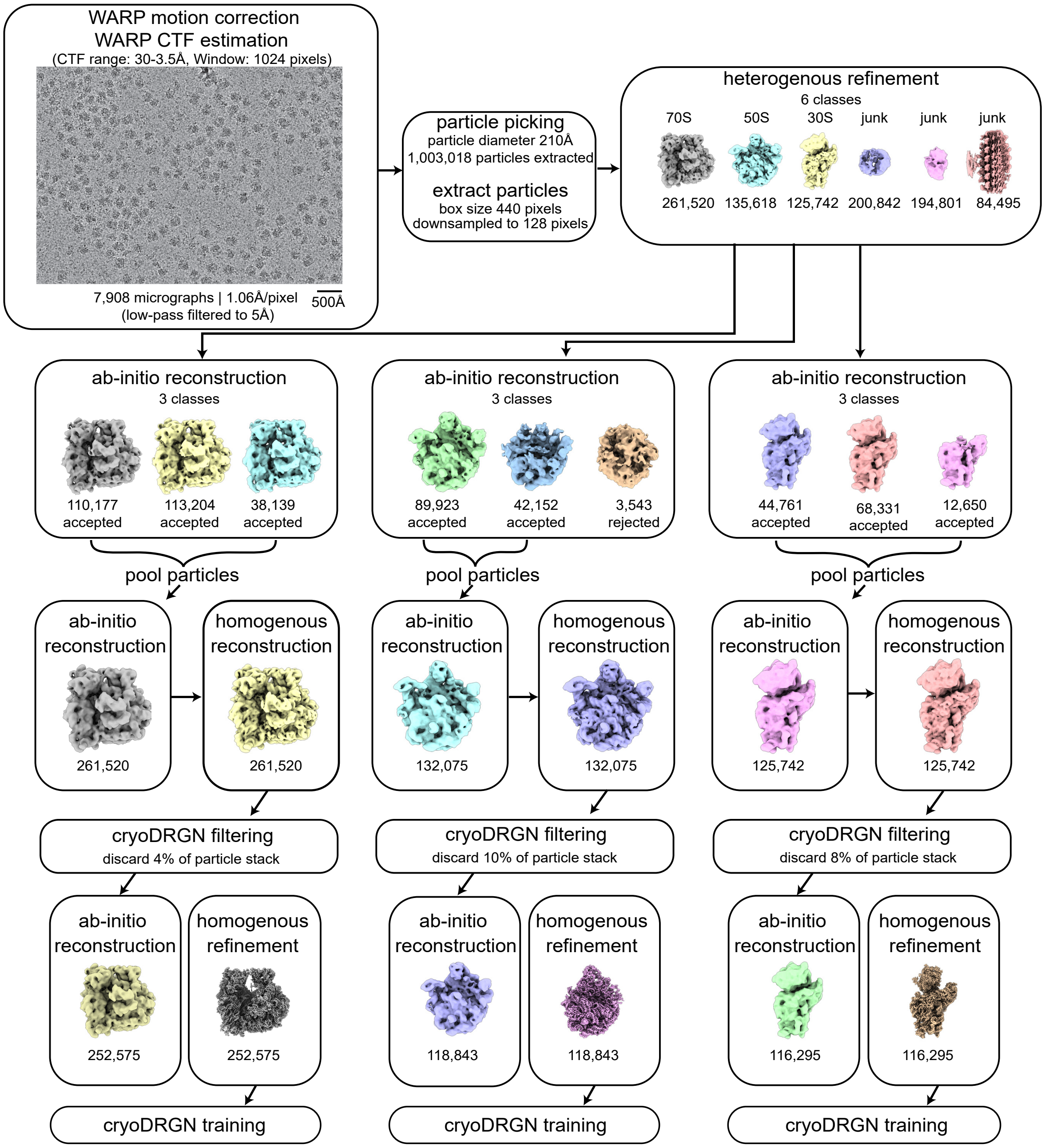


**Figure S12. CryoSPARC pre-processing workflow for analysis of the 25°C dataset.**
The job names and key parameters are listed, with particle numbers listed and representative maps depicted.

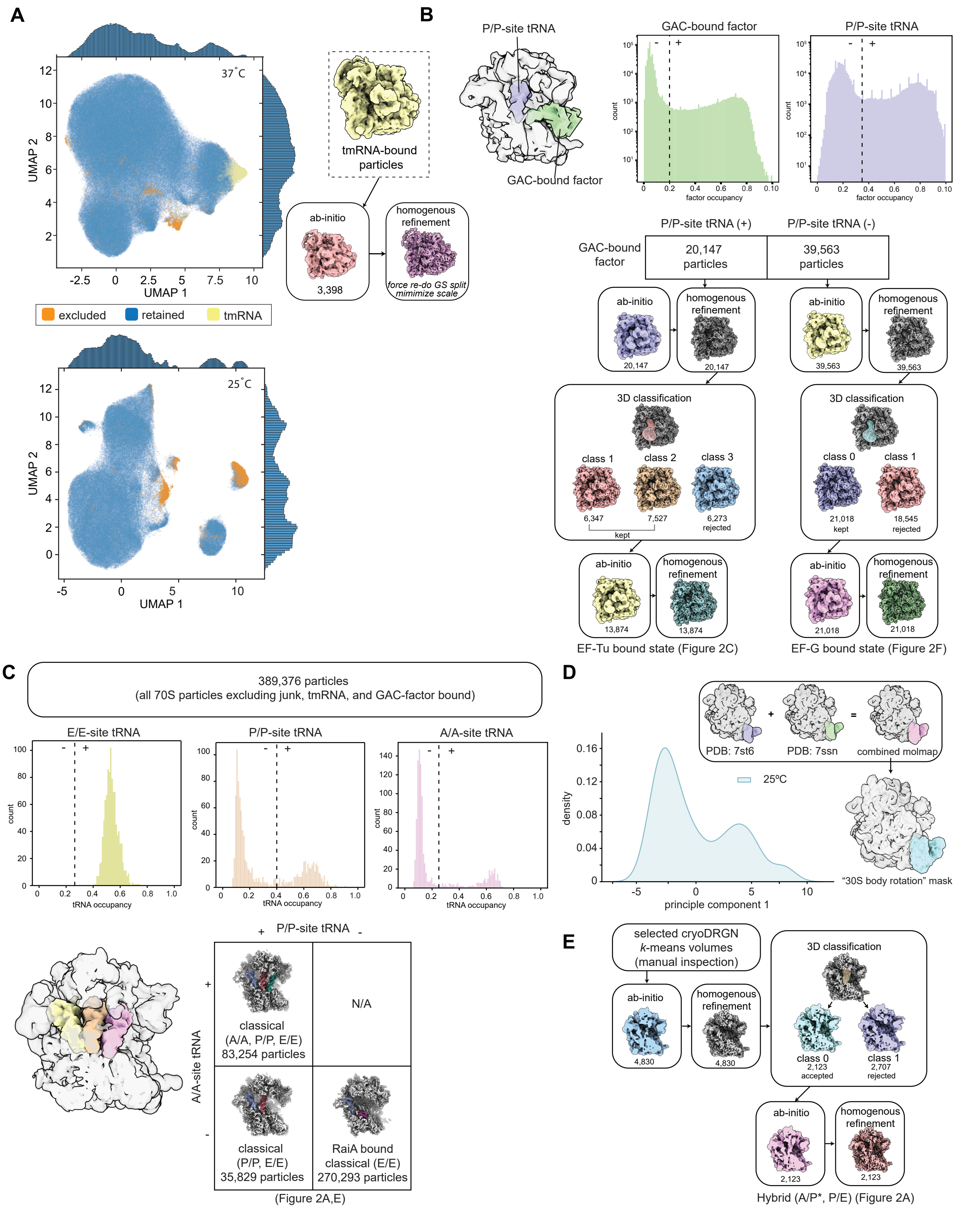


**Figure S13. Image processing workflow for analysis of 70S ribosome structural states.**

**(A)** Scatter plot of cryoDRGN latent space for all 70S ribosomes after UMAP dimensionality reduction. Each point marks a projected particle from the 37°C (top) or 25°C (bottom) dataset. Marginal distributions plotted. Particles were filtered by magnitude of the cryoDRGN 8-dimensional latent variable to exclude poorly posed or undesired particles (orange). The tmRNA-bound particles (yellow), as identified by inspection of *k*=50 *k*-means centroid volumes clustered in this space, with a representative volume depicted in yellow. These particles were then used in cryoSPARC ab-initio reconstruction and homogenous refinement, with each resulting volume depicted. *(continued)*

**Figure S13 (*continued from prior page*).**

**(B)** Processing workflow for EF-Tu and EF-G-bound ribosomal states. CryoDRGN volumes were generated for each particle at box size 64 and MAVEn '*on-the-fly*' occupancy analysis was performed within depicted masks for P/P-site tRNA (purple; derived from 7st6 Carbone *et al.* 2021) and a GTPase associating center (GAC) (green; derived from 6wd2 Loveland *et al.* 2020), which are overlayed on a representative cryoDRGN map reconstructed at box size 64 (grey). Resulting occupancy for these elements is plotted as histogram. Dotted lines depict thresholds defining presence or absence of P-site tRNA and GAC-bound factor. Volumes bearing density in the GAC were further classified based on the presence or absence of P/P-site tRNA, with particle counts noted in table. Each resulting particle stack was exported to cryoSPARC and ab-initio reconstructions were used as input models for homogenous refinements, with resulting maps depicted. 3D-classification with a focused mask around the GAC was then applied, revealing classes with density corresponding to either EF-Tu (workflow on left) or EF-G (workflow on right). These particles were pooled and used for subsequent ab-initio and homogenous refinement, resulting in reconstructions shown in Figure 2C and Figure 2F, respectively.

**(C)** Classification of ribosomes with tRNAs in their classical 'A/A, P/P, E/E' configuration. Using particles remaining after excluding those analyzed in (A) and (B), 1,000 cryoDRGN volumes were sampled from *k*=1000 *k*-means centroid locations and were analyzed using MAVEn occupancy analysis within masks derived from 7st6 for the E/E, P/P, or A/A-site tRNAs, which are overlayed on a representative cryoDRGN volume in yellow, orange, and pink, respectively. Resulting occupancy for each mask plotted as a histogram with thresholds used to distinguish presence (+) vs absence (-) of each tRNA marked with a dotted line. All particles bore E/E-site density. Particles stacks as listed in the contingency table were each exported to cryoSPARC and refined using homogenous refinements with per-particle defocus refinement enabled, with resulting reconstructions depicted. Inspection of these maps revealed that particles lacking A/A- and P/P-site tRNA contained RaiA.

**(D)** Voxel principal component analysis (vPCA) from Figure 2B. A large mask spanning 16S rRNA helices 6, 10, and 17 in both the non-rotated and hybrid states (derived from 7st6 and 7ssn, respectively) was generated using the scheme shown.

**(E)** Classification of hybrid tRNA states (A/P*, P/E) at 25°C. CryoDRGN volumes were sampled from *k*=200 *k*-means centroid locations of the full 70S particle stack and volumes with hybrid tRNA density were identified by manual inspection of the volumes, resulting in 4,830 particles. These particles were used as input for ab-initio reconstruction, homogenous refinement, and 3D-classification using a mask around the A/P*-site tRNA (as defined in 7ssn and depicted in brown) in cryoSPARC. Particles in class 0 were used for a second round of ab-initio reconstruction and homogenous refinement, resulting in the reconstruction shown in Figure 2A.

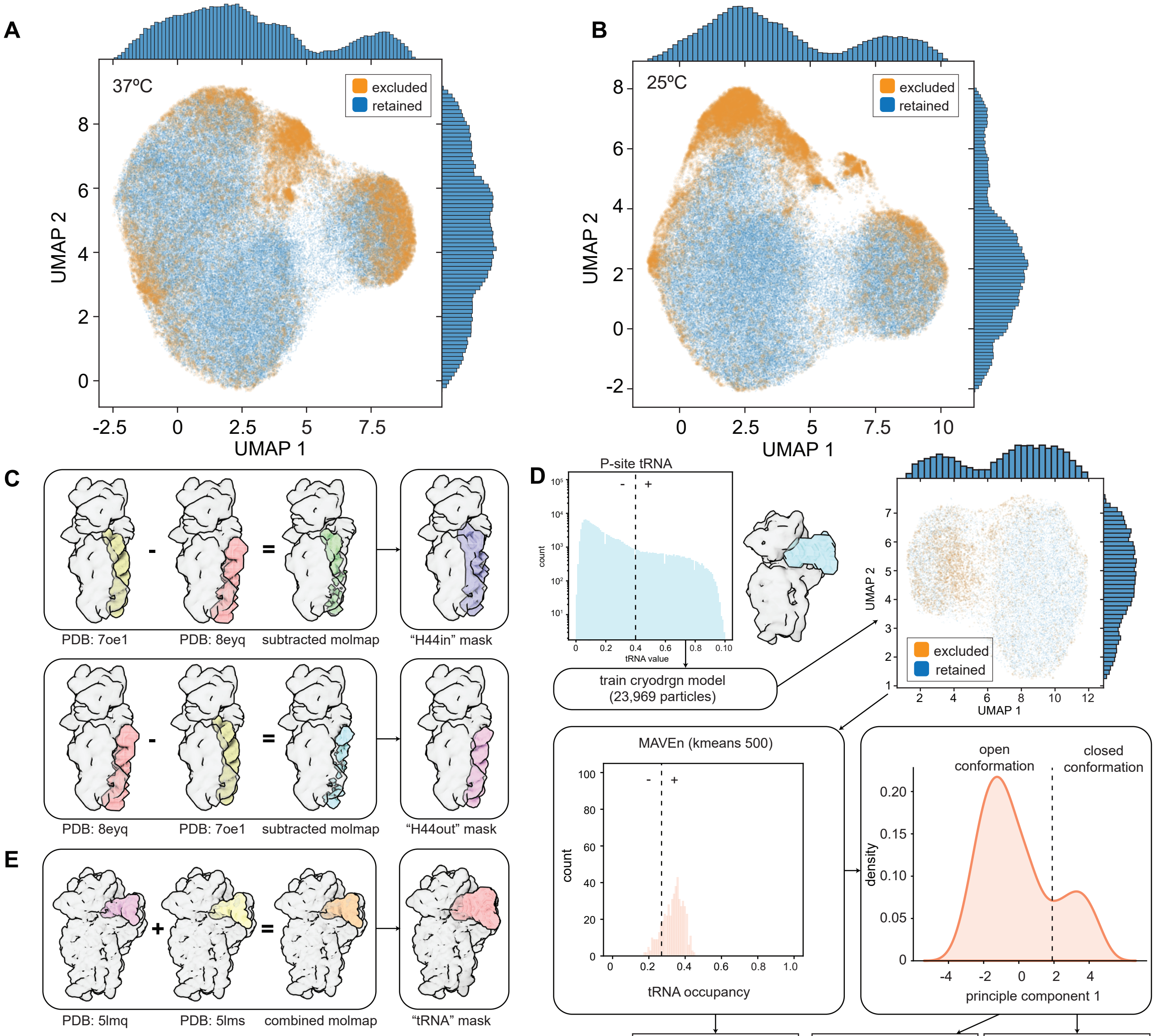


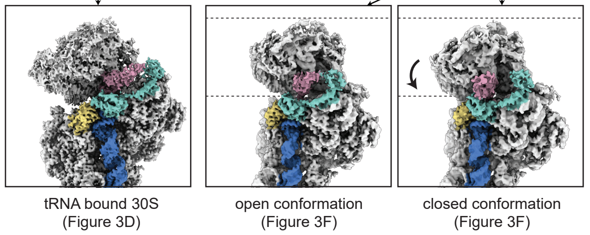


**Figure S14. Image processing workflow for 30S states.**

**(A-B)** Scatter plot of cryoDRGN latent space for all 30S subunits after UMAP dimensionality reduction. Each point marks a projected particle from the 37°C (A) or 25°C (B) dataset. Particles were filtered by magnitude of the latent variable to exclude poorly posed or undesired particles (orange).

**(C)** Mask generation scheme for H44 classification. Density maps for mature H44 and dislodged H44 were generated from PDBs 7oe1 (Maksimova *et al.* 2021) and 8eyq (Sun *et al.* 2023), respectively, using the ChimeraX `molmap` command. Density maps were subtracted as shown and converted to masks as described in the methods.

**(D)** Classification of P-site tRNA-bound 30S subunits. Histogram of P-site tRNA occupancy as calculated by MAVEn '*on-the-fly*' analysis using a mask (blue), which is depicted overlaid on representative cryoDRGN volume. Dotted line denotes occupancy threshold used to extract particles (n=23,969) with high occupancy within mask. These particles were used to train a cryoDRGN model, and were filtered based on magnitude of the latent embedding vector and manual inspection of *k*=50 k-means centroid volumes. Retained particles (n=19,195) were used for a subsequent round of cryoDRGN training, and 500 volumes were sampled from *k*=500 k-mean centroid locations and analyzed using MAVEn occupancy analysis with a "tRNA" mask, as defined in (E), and with resulting occupancy plotted as a histogram. Particles with a fractional occupancy greater than 0.27 (noted with dotted line) were used to generate the homogenous refinement for tRNA bound 30S (Figure 3D). The same particles were used as input for voxel principal component analysis (Sun *et al.* 2023), with the resulting distribution plotted. Particles below a threshold of 1.9 along the first principal component were used as input for a homogenous refinement in the "open conformation" and particles above a threshold of 1.9 were used as input for a homogenous refinement in the "closed conformation", as displayed in Figure 3E.

**(E)** Mask generation scheme for tRNA. Density maps for two unique tRNA conformations were generated from 5lmq and 5lms (Hussain *et al.* 2016), summed, and converted to masks as described in the methods. Masks are overlaid on a single 30S cryoDRGN volume.

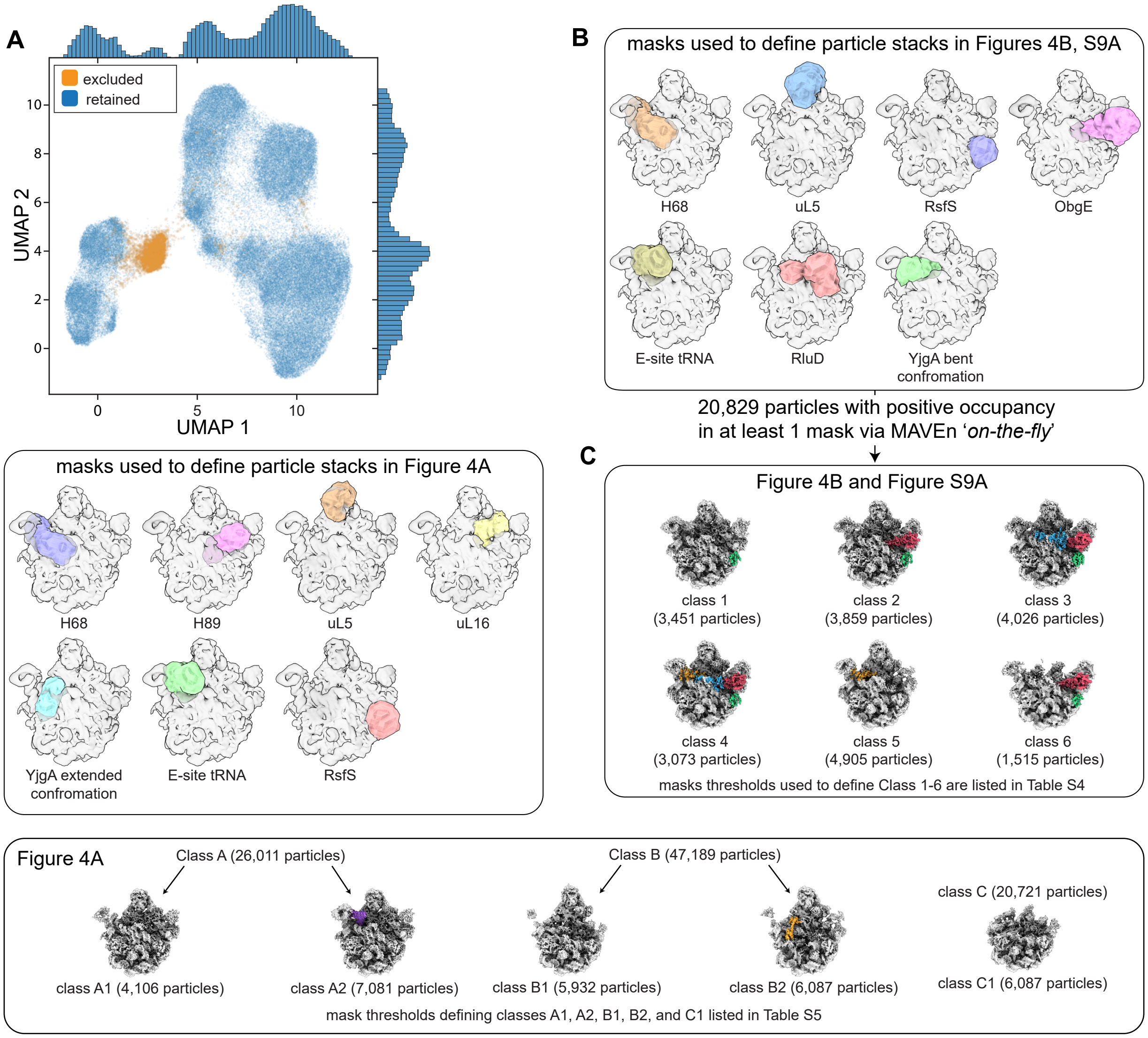


**Figure S15. Image processing workflow for heterogenous 50S states.**

**(A)** Scatter plot of cryoDRGN latent space for all 50S subunits after UMAP dimensionality reduction. Each point marks a projected particle from the 25°C dataset. Particles were filtered using volumes sampled from *k*-means (*k*=50) centroid locations to remove clusters of poorly resolved particles (orange).

**(B)** Depiction of masks (colored) used in the MAVEn 'on-the-fly' analysis to isolate particle stacks used in cryoSPARC homogeneous reconstructions of factor-bound 50S particles, as described in Methods. A single 50S cryoDRGN volume (box size 64) is shown in translucent grey for reference. Table S4 notes per-mask occupancy thresholds used to define each class.

**(C)** Density maps resulting from homogeneous refinement of particle stacks isolated using masks in (B), with number of particles noted. Maps are colored using docked atomic model for bound factors RsfS (green), YjgA (yellow), RluD (red), and ObgE (blue).

**(D)** Depiction of masks (colored) used in the MAVEn 'on-the-fly' analysis to isolate particle stacks used in cryoSPARC homogeneous reconstructions of 50S assembly states, as described in Methods. A single 50S cryoDRGN volume (box size 64) is shown in translucent grey for reference. Table S5 notes per-mask occupancy thresholds used to define each class.

**(E)** Density maps and particle count resulting from homogeneous refinement of particle stacks isolated as follows. First, MAVEn occupancy analysis was used to partition the 93,921 particles that were not analyzed in (B-C) as described in Methods, and depicted in Figure S6. These particles were then subjected to MAVEn '*on-the-fly*' analysis using the masks from (D), and particle subsets as defined by thresholds in Table S5 were isolated and refined using cryoSPARC.
